# Mechanical response of the RecA nucleoprotein filament to increasing D-loop length

**DOI:** 10.64898/2026.02.03.702038

**Authors:** Afra Sabei, Alexandre Détruit, Sébastien Neukirch, Claudia Danilowicz, Mara Prentiss, Chantal Prévost

## Abstract

Protein filaments play fundamental functions in the cell, ranging from scaffolding like in the cytoskeleton to sensing and transmitting forces and torques. Here we address the case of the nucleoprotein filaments (NPFs) of homologous recombination (HR) formed by the polymerization of the RecA protein on DNA. In contrast to the cytoskeleton filaments, the HR filaments are not known to exert or sense forces. However the stress in the stretched and unwound DNA bound to those filaments was shown to play a role in promoting DNA strand exchange during the early stage of the HR mechanism. Here we use molecular dynamics simulations to examine whether the strain in the nucleoprotein filament upon strand exchange progression and D-loop formation may influence subsequent steps of the HR process. Our results indicate that the filament mechanical properties are sensitive to the length of DNA incorporated in the D-loop. The response we observe upon increasing the D-loop length is first elastic, up to a threshold that we estimate to be 27 incorporated base pairs, after which the NPF enters a new stage where the protein-DNA connectivities are reorganized. Notably, the DNA displaced strand locally switches from site II to site III, a newly characterized binding site. We discuss the possible consequence of this mechanical response of the NPFs for the course of the HR process.

## 1 Introduction

Filaments commonly found in the cells exert essential functions such as cell wall support, network for the transport of vesicles, chromosome segregation or partitioning [1], but also force generation and force sensing [2], [3]. Force generation most often relies on polymerization/depolymerization at the filament extremities. It is used for example by microtubules to separate the chromosomes during meiosis [4], [5]. Force sensing has been less studied but a growing body of observation suggests that exerting a longitudinal force or a torque on actin filaments modulates the association rate of actin-binding proteins [6]. More generally, protein filaments have been proposed to transmit forces in a collaborative way via chains, membranes or other filaments they use as scaffolds [7].

Recombination nucleoprotein filaments (NPFs) are right-handed helical filaments that assemble on DNA single strands resulting from DNA double strand break damage. Like many other protein filaments, NPFs control their length by polymerizing (preferentially in 3’ of the bound DNA) and depolymerizing (preferentially in 5’). Unlike for cystoskeleton filaments, the polymerization/depolymerization process does not seem to be associated to force production, such as pulling or pushing. However as discussed below, experimental data reported by our team and others [8], [9], [10] suggest NPFs may sense forces and respond to them. The present work aims at investigating and characterizing this potential response in a structural and dynamical point of view.

Homologous recombination faithfully repairs DNA double-strand breaks (DSBs) by using homologous genomic double-stranded DNA (dsDNA) as a template [11], [12], [13], [14], [15]. To this aim, dsDNA regions are incorporated in recombination nucleoprotein filaments formed on DNA single strands (ssDNA) that result from DSB processing. The dsDNA sequence is probed via base pairing exchange between the ssDNA and the dsDNA. If the sequences match, the ssDNA captures the dsDNA complementary strand (a process called strand exchange), leading to a RecA-bound heteroduplex DNA product and a displaced strand, the three strands being referred to as a D-loop. The initial recognition stage is limited to 8 base pairs and is extremely quick, in the order of *µ*s, enabling multiple trials to be performed in a short time. When the test is passed, incorporation of new DNA bases occurs by successfully probing groups of three bases (or triplets) [16], in a much slower kinetic regime (0.5 s per incorporated triplet on average). In the absence of ATP hydrolysis and for lengths of incorporated DNA greater than 20 to 30 base pairs, roughly corresponding to one and a half filament turns, the strands remain firmly bound inside the filament. For shorter lengths of incorporated DNA, the heteroduplex product is unstable regardless of whether ATP is hydrolyzed or not, and the probability of product reversal is very high [8]. Finally, in the presence of ATP hydrolysis, the filament-bound DNA product rarely extends beyond 50 to 80 base pairs, as shown both *in vitro* [8], [17] *and in vivo* [9], [10]. Experimental evidence shows that above these distances, the filament unbinds from the heteroduplex product in 5’ of the exchanged region [8], [17]. In addition, we previously showed that the final stage of the HR mechanism, *i.e*. the passage to DNA synthesis by a DNA polymerase, also depends on the length of incorporated and exchanged dsDNA, with the efficiency of DNA synthesis increasing for lengths greater than 50 base pairs [18]. Altogether, these results suggest a kinetic control that would depend both on ATP hydrolysis and on the length of incorporated DNA. In this work, we focus on the part of the control process that specifically depends on the length of incorporated DNA.

In NPFs, the DNA strands are stretched by 50% and unwound by 40%[19]. The fact that the searched dsDNA presents different levels of extension inside or outside the filament has fed enlightening discussion about the efficiency of sequence recognition [20], [21]. Further work showed that strand extension also plays a role in the mechanism of strand exchange itself [22], [23]. One factor comes from the differential extension of the DNA strands once in the filament. The primary DNA binding site (site I) that first binds the ssDNA in the pre-synaptic complex and then binds the heteroduplex resulting from strand exchange in the D-loop (post-synaptic complex) is situated close to the filament axis [24]. In contrast, the secondary DNA binding site (site II) that binds the dsDNA and the displaced strand respectively in the synaptic and post-synaptic complexes is laterally distant from the axis, as anticipated by E. Egelman back in the 90’s [25]. This results in the DNA strand backbones being more extended in site II than in site I (see Figure 4 in [23]). Using single molecule experiments, the Prentiss group showed that the tension that differently builds in the DNA strands is a strong driving force for strand exchange [22], [23], [26]. In addition, simulations showed that in the absence of strand exchange, the filament site II cannot accommodate more than 12 dsDNA base pairs; incorporating additional groups of three dsDNA base pairs [27], [28] results in sudden dissociation of the dsDNA from site II [23], a result that was recently confirmed by Cryo-EM observations [29]. As a whole, the various dependencies to the length of exchanged DNA suggest that the stress in incorporated DNA may influence the mechanical and dynamical properties of the whole macromolecular assembly.

Physics tells us that the response to stress in any material, called the strain, is primarily an elastic response characterized by quantities such as the stretching and torque moduli; after a certain threshold has been reached, the elastic response gives way to a plastic state characterized by internal reorganization of the material under tension. In the case of the RecA NPF bound to a D-loop, an elastic response would be expected to take the form of global distortions in the filament in terms of bending, twisting or stretching, while a plastic response would involve reorganization of the network of interactions in monomer-monomer interfaces or protein-DNA contacts. If it can be established that the NPF properties vary with successive incorporation of dsDNA triplets, then this will indicate that stress is building and the response can be interpreted in terms of the filament mechanics. Understanding how the nucleoprotein assembly made of non-covalently bound subunits responds to the tension in the DNA strands may then be a clue to understanding the length-dependency observations in the recombination process.

Obtaining such information via experimental methods, *e*.*g*. by capturing intermediates or via single molecule manipulation, remains highly challenging. Single molecule experiments have brought huge progress in the understanding of the HR process [26], [30], [31] but in the present case, they may be hampered by the requirement that control must be exerted on at least one DNA strand, which might interfere with forces or torques inside the filament and may bias the process. In these conditions, numerical simulation appears as a promising method to explore possible response to perturbations in the HR filaments.

In this study, we use molecular dynamics simulations to investigate the NPF response to successive incorporation of dsDNA triplets. We address the question of whether the filaments are sensitive to the tension of the bound DNA strands and if so, how they adapt to that tension. To this aim, we built models of the post-synaptic filament where the D-loop length varies between 9 and 54 base pairs (bp) by groups of 3 base pairs and we investigated the short-time response of the filaments to added DNA in terms of global distortions in the protein filament and perturbations of the protein-DNA contact network. This leads us to identify a length threshold of 27 base pairs where the elastic response gives way to a plastic response. We observe that in the plastic regime, the displaced strand partly migrates to a newly characterized DNA binding site III and we discuss the possible consequences for the homologous recombination mechanism.

## 2 Material and Methods

### 2.1 Construction of NPFs with varying D-loop sizes

The construction started from the CryoEM structure with PDB code 7jy9 [29]. That structure is composed of 9 RecA monomers bound to a 27-nucleotide single-stranded oligonucleotide (ssDNA) and a 42-base pair (bp) double-stranded DNA (dsDNA). In the original structure, the 42-bp dsDNA presents a 10 bp-long heterologous bubble starting at position 16, designed to artificially stabilize a 10 bp-long D-loop resulting from local annealing of the ssDNA and the dsDNA complementary strand into a heteroduplex **(Figure 1A)**. We first modified the sequence of the CryoEM structure leading strand (also called displaced strand in this paper) so that the homology with the complementary strand was restored over the whole dsDNA length (see the sequence in **Figure 1C**). This makes reverse base pairing exchange possible in the D-loop and indeed, the last base pair in the 3’ end of the heteroduplex region was found to be unstable during molecular dynamics simulations. We refer here to the modified CryoEM structure as the 9bp model, after the number of stable heteroduplex base pairs.

**Figure 1:**
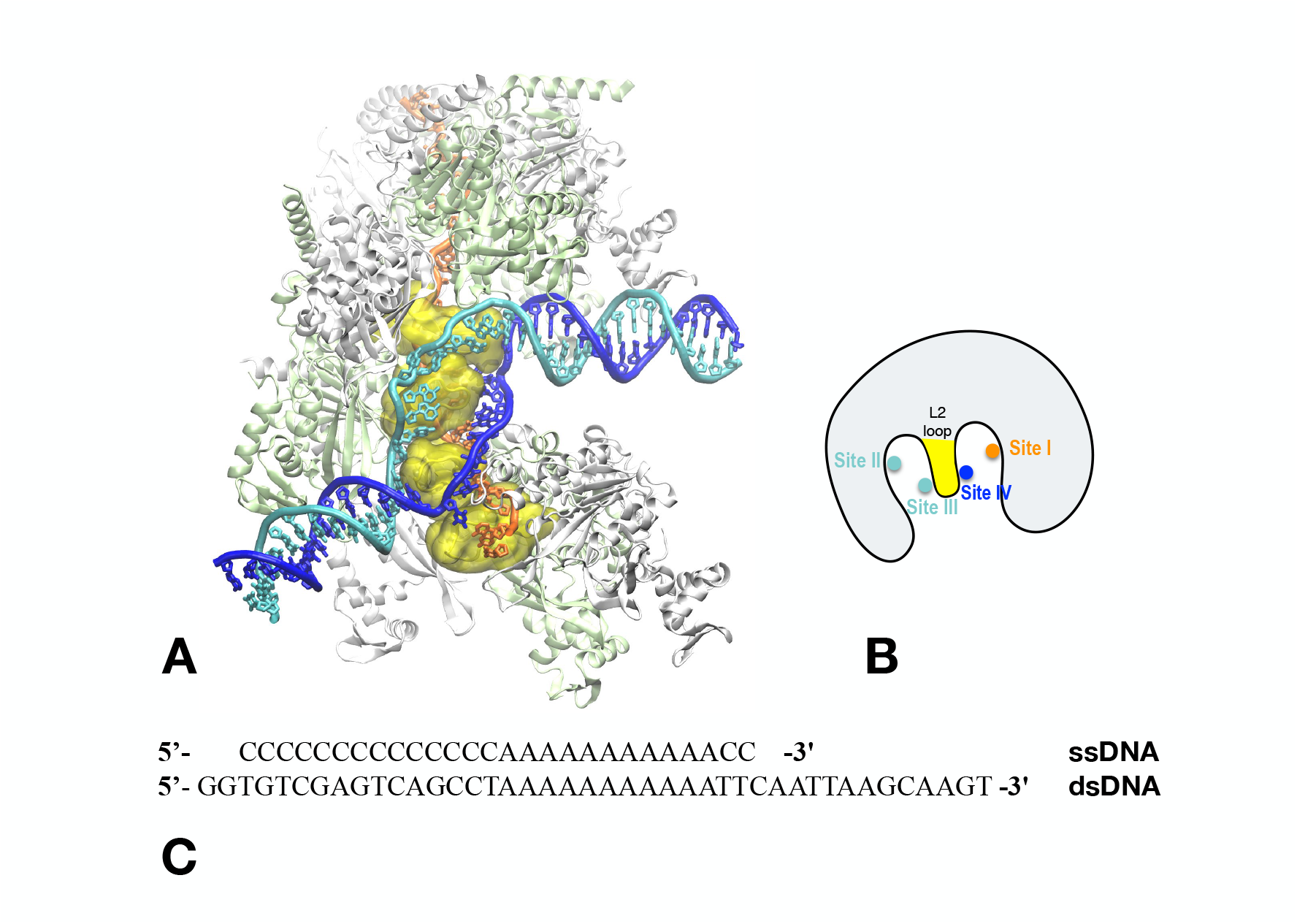
The 9bp model. (A) Graphical representation of the structure; protein monomers are represented in white and lime alternatively; the DNA strands are in orange (ssDNA), blue (complementary strand) and cyan (displaced strand); the bulky L2 loops that separate the outgoing and complementary strands in the D-loop are shown in yellow, surface representation. (B) Schematic representation of a filament slice taken perpendicularly to the axis, showing the DNA binding sites at the beginning of the MD simulations (site I, site II, site IV) and an alternative site for the displaced strand from the longest D-loop sequences (site III); the colors for the DNA strand positions are the same as in (A). (C) Sequence of the DNA strands. The VMD software [44] is used for all graphical representations in Figures 1 to 5.

We subsequently generated models with increasing length of the D-loop region. The construction consisted in replicating a subunit formed by the central monomer and the three DNA triplets it binds to. We applied to this subunit the transformation that superposes the monomer quasi-rigid core region (excluding the L1 and L2 loops and the N- and C-terminal domains) to the same region of its neighbor monomer in the 3’ direction using the module Heligeom of the PTools library [32], [33] (an example of step by step building protocol using similar principle is described in reference [34]). This procedure produced a site I-bound heteroduplex where the two backbones were almost perfectly continuous; in contrast, careful minimization using the NAMD software [35], [36] (versions 2.12 and 2.13) was necessary to close the backbone of the displaced strand in site II. Site II is laterally displaced with respect to the axis, which increases the stretching of the displaced strand backbone with respect to the two strands in site I. During minimization, the phosphate atoms on each side of the strand disruption locations every three bases were initially restrained to their initial position. The models are called after the number of stable heteroduplex base pairs, *i*.*e*. the 12bp model results from duplicating the central subunit, the 15bp model from triplicating this subunit, the (6+3n)bp model from replicating n times the subunit. The 12bp model differs from the other models by the presence of an additional ss-DNA bound 3-monomer filament segment in 3’ of the D-loop-bound region; this enables exploring the possibility for the B-form dsDNA in 3’ to make contacts across the filament groove, which cannot be observed if the filament ends immediately in 3’ the D-loop like in the CryoEM structure. For the 21bp system, a model without the B-form terminal regions was derived from the full model. Finally, a 36bp simulation is taken from previous modeling work where protein DinB was bound to the filament extremity in 3’ [37]; contacts in the filament extremities are therefore not exploitable for that simulation, however the fact that contacts in the center of the D-loop present trends analogous to what is observed for the 54bp prompted us to display these results as supplementary information.

### 2.2 All-atom Molecular Dynamics (MD) simulations

#### Simulation setup

All systems were prepared using VMD auto PSF builder(VMD 1.9.2).The structures were first solvated using a TIP3P water model in boxes submitted to periodic boundary conditions. We used a physiological ionic concentration of 0.15 mol/l (see Table 1 for a detailed composition of the simulated systems). Overall the simulated systems contained between 435,000 and 900,000 atoms. Molecular dynamics (MD) simulations were performed on all systems using the NAMD software[36], versions 2.12 and 2.13, and the CHARMM36m force field including the CMAP correction [38] under periodic boundary conditions with the Particle Mesh Ewald (PME) method [39]. Time steps were set to 2 fs using the SHAKE algorithm [40]. Temperature and pressure were regulated via a Langevin dynamics approach, employing a Nosé-Hoover-Langevin piston for control. Throughout the equilibration phase, the protein C*α* carbon and DNA P positions were harmonically restrained to their initial coordinates, with the force constant reduced incrementally from 0.5 to 0.05 kcal·mol^−1^·Å^−2^ during 60 ns. During the 200 ns production phase, no restraints were applied. All simulations were replicated, so that altogether we have 2*200 ns of production for each complex.

**Table 1:** Summary of the MD simulations performed in the study (for each system, we ran two replicas of 200 ns).

| System | Protein Atoms | Monomer | ssDNA | dsDNA | Nucleic atoms | Na<sup>+</sup> Ions | Cl<sup>-</sup> Ions | Water Molecules |
| --- | --- | --- | --- | --- | --- | --- | --- | --- |
| 9bp | 42,771 | 9 | 27 | 41 | 3,503 | 433 | 288 | 101,936 |
| 12bp | 62,517 | 12 | 38 | 51 | 4,473 | 609 | 419 | 148,336 |
| 15bp | 48,090 | 10 | 29 | 48 | 3,996 | 503 | 340 | 120,519 |
| 18bp | 57,198 | 11 | 36 | 51 | 4,374 | 543 | 359 | 128,945 |
| 21bp | 57,708 | 12 | 39 | 54 | 4,662 | 582 | 390 | 138,082 |
| 27bp | 71,625 | 15 | 45 | 56 | 5,238 | 649 | 426 | 150,630 |
| 45bp | 106,296 | 21 | 63 | 78 | 7,569 | 886 | 585 | 207,163 |
| 54bp | 110,607 | 24 | 68 | 87 | 7,740 | 985 | 653 | 231,286 |

#### Analysis

All trajectories were initially aligned on the single stranded DNA. The analysis of the molecular dynamics trajectories was conducted using VMD analysis tools and custom Python scripts, employing the MDAnalysis [41], [42] and MDTraj libraries [43].

### Evolution of the distances between DNA strands and their binding sites during molecular dynamics simulations

For each system, the trajectory files were processed using the MDAnalysis Python library (version 2.9.0). Distances between the displaced strand and site II on the one hand, the ssDNA and site I on the other hand, were computed based on selected groups of atoms in each of the nucleic or protein partner. For the displaced strand, the selected group included phosphate group atoms (phosphorus atom P and bound oxygen atoms O1P, O2P, O5’, O3’) from a specified range of nucleotide residues corresponding to the D-loop region starting from base 19 in all cases. Site II of the protein was identified by the clusters of basic residues implicated in DNA binding, *i*.*e*. Arg226, Arg227, Lys232, Arg243, Lys245; site I was defined by the L1 loop residues 161-164 and Arg169. For each frame in the trajectory, the minimum distance between the two groups was computed using pairwise distance matrices, on a per-residue basis. For the analysis shown in **Section 3.3**,, the per-residue minimum distances were averaged to yield a single distance value per frame, representing the proximity of the displaced strand to site II over time. Details of the per-residue time evolution have also been computed and are displayed in supplementary information.

### Contact map calculations

In this analysis we focused on bases within the displaced strand of the nucleic acid. For each DNA base, the proximity to protein residues across multiple protein chains was examined. At each simulation frame, the minimum distances between all atoms of the selected DNA residue and protein residues were computed, and the percentage of frames where these distances were below a 5 Å cutoff was recorded. This metric quantifies the frequency of close contacts between DNA and protein residues throughout the trajectory, providing insight into the stability and nature of the DNA-protein interface.

### f_NAT_analysis

In this analysis we used the fnat.py script from the PTools library [32]. Sets of contacting residue pairs between adjacent monomers are extracted from the starting model, taken as the reference. Residue pairs are considered top be in contact if any atomic pair from each residue across the interface is within a specified cutoff distance of 5 Å. The initial contacts are then compared with those observed in each MD frame and the fraction of preserved contact in the frame is computed.

This metric reflects how similar the interfaces are, with higher f_*NAT*_ values indicating closer preservation of the original contacts. It is adapted from the metric used for the evaluation of docking simulation [45], where the reference structure is the native structure hence the name f_*NAT*_ (fraction of native contact).

### Filament axis

Axis were generated either as simple broken lines or as elastic rods, following the principle described below. For each pair of interacting RecA monomers P1 and P2, we computed the screw transformation that leads from from P1 to P2 rigid parts (core domain without loops L1 and L2) [33], [46]. The screw transformation is characterized by an axis, a rotation around this axis and a translation along the axis. An axis segment was defined based on these characteristics, centered on *M*, the projection on the axis of the middle point *C*_*M*_ between the centers of mass *C*_*M*1_ and *C*_*M*2_ of P1 and P2 and with length the translation value as reported (see supplementary information **S1, Scheme S1A)**; concatenating the axis segments relative to successive pairs of interacting monomers produces a broken line which was used to compute the end-to-end distance *EE*, the contour length *L* and the axis shortening parameter *AS* such that *AS* = 1 − *EE/L*. The contour length *L* is measured on the starting model, which is a straight helix. The broken line is discontinuous and does not convey the torsional information present in the screw transformation parameters. We therefore defined reference frames that were used to generate elastic rods following the method described in [47]. The reference frames were positioned at the center *M* of each axis segment, with unit vectors **u** in the axis direction, **v** pointing towards point *C*_*M*_, and **w** as the cross product between **u** and **v** (SI1, Scheme S1B). The elastic rod was constructed from segments that link one frame to the next one in a continuous way relative to position and torsion parameters. Finally, the axis points M defined above were used to compute local curvature LCs and flexibility LFs using the Menger-based metrics reported in [48] and available as a MDAKit from the MDAnalysis package [49]. That metrics showed very useful in characterizing the structural properties of intrinsically disordered chains as well as folded chains. Here, we find it also can reproducibly capture dynamic trends in the filament axes deviation from linearity, in spite of the short simulation times. All the scripts can be found in the zenodo repository https://zenodo.org/records/18377475, along with the MD trajectories.

## 3 Results

This work aims at getting insight into the structural and mechanical effects of incorporating an increasing number of base triplets into RecA NPFs undergoing strand exchange. For this, we constructed post-synaptic filaments with different D-loop lengths and we submitted these models to molecular dynamics (MD) simulations. The construction started from the CryoEM structure with PDB code 7jy9 [29], after modifying the sequence of the dsDNA leading strand (also called displaced strand) to regain sequence complementarity (see Material and Methods and **Figure 1A,C)**. Models with successively added groups of three base pairs correspond to intermediate states in the D-loop elongation process, as identified in our earlier simulation work [23] and recently confirmed both for prokaryotic RecA [50] and eukaryotic Rad51 [51], [52], [53]. They are referred to after the number of stable heteroduplex base pairs in the D-loop (e.g. 9bp, 12bp, …)

It is important to note that due to the construction protocol, added RecA-DNA regions present uniform geometries at the beginning of the molecular dynamics (MD) simulations. The simulations are short and cannot be expected to provide information on the long-term evolution of the system or its thermodynamic equilibrium, but this is not the object of the study. What we aim to examine is the immediate response of the system to perturbation resulting from the incorporation of new DNA triplets: deviations from the initially uniform geometry will inform about the presence of tension in the system and if so, how this tension gets dissipated. We repeated the simulations to improve the reliability of the observations. **Table 1** recapitulates the models that were submitted to MD simulations. In this section, we analyze the MD trajectories by separately characterizing the global properties of the filament, the network of protein-DNA and protein-protein interactions, and examining how these characteristics evolve with the D-loop length.

### 3.1 An extended network of protein-protein and protein-DNA interactions

RecA nucleoprotein filaments result from the non-covalent assembly of RecA proteins on DNA strands. Using single molecule manipulation, the group of Bustamante showed these assemblies are globally very stiff in the absence of ATP hydrolysis, with a persistence length of about 950 nm (close to 10 filament turns) for filaments constructed on either single-stranded or double-stranded DNA [54]. As highlighted in that study, the stiffness arises in large part from the network of interactions between the protein monomers and the DNA strands. The main protein-DNA interactions sites have long been characterized [14], [24], [55]: the DNA binding site I, that binds the damaged single-stranded DNA alone or as part of the heteroduplex resulting from strand exchange, is a narrow cleft situated between the flexible loops L1 (residues 156-164) and L2 (194-212) (**Figure 1A,B**); the DNA binding site II, that binds the displaced strand in the synaptic complex [23] or after strand exchange [23], [24], [29] is also a narrow cleft located between loop L2 and a pending hairpin in the core domain sometimes referred to as LexA-binding loop and that we will call here the S2 hairpin (S2 for site II; residues 225-245). Interestingly, the L1, L2 and S2 motifs belong to different monomers: if the central L2 loop belongs to monomer (*i*), then the L1 loop associated with L2 to define site I belongs to monomer (*i* + 1) while the S2 hairpin that contributes along with L2 to defining site II belongs to monomer (*i* − 1). This means that binding a three-strand triplet level of a D-loop involves three successive monomers forming intertwined interactions. Both site I and site II are strongly electropositive, but the electrostatic field is stronger for site II which contains a cluster of basic residues; the field remains strong in site II even when a DNA strand occupies site I (supplementary Figure S2 in [28]). The complementary strand can occupy two additional sites that we will call sites III and site IV **(Figure 1B and supplementary Figure S2)** – and indeed, former experimental studies point out the possibility for the RecA filament to simultaneously accommodate four nucleotidic strands [56], [57]. Our former simulations of the initial pairing exchange process argue for the complementary strand transiently occupying site III in the synaptic state before strand exchange takes place and results in its migration to site IV [23]. Site III and site IV are both situated on the L2 loop surfaces of successive monomers, but site III is on the same side of the L2 loops as site II, while site IV is on the same side of L2 as site I. The complementary strand does not bind strongly to either of these binding sites, which mostly contain hydrophobic residues.

At the level of the filaments, the network of interactions in the CryoEM structure between individual protein monomers and DNA strands is dominated by the protein-DNA interactions as can be seen from interaction maps generated by the Mapiya website [58] (supplementary **Figure S3A**). In that structure, each protein monomer interacts mostly with its two neighbor proteins and shows limited contacts with its two next-neighbors. However, the ssDNA strand contacts all protein monomers and the dsDNA strands contact most monomers. **Figure S3B** also shows that even for the stable 9bp structure, new interactions can form during MD simulations with notably new protein-DNA or protein-protein interactions appearing across the filament groove.

### 3.2 Increasing the length of incorporated DNA modulates the filament mechanical properties

In the absence of applied stress, the filaments naturally undergo structural fluctuations due to thermal motions that take place in protein monomers, DNA strands, protein-protein and protein-DNA interfaces [59], [60]. If additional stress is transmitted to the filament by strained DNA strands, a possible response can primarily be collective and will then concern the global dynamic behavior of the filament. We first examined how the filament responds to the presence of increasing D-loop lengths.

To this aim, we defined an axis for the filament from successive screw transformations that relate each monomer to the next one in the filament (see Methods and **Supplementary information S1**). The axis is either defined as a succession of axis segments (**Scheme S1A**), an elastic rod (**Scheme S1B**) or a curvilinear axis (**Scheme S1C**). To describe the overall deformation, we first considered the Axis Shortening metrics *AS*, defined from the ratio between the end-to-end distance (*EE*) and the contour length (*L*) of the filament axis, *AS* = 1 − *EE/L*. **Figure S4** shows that the information provided by this simple descriptor, averaged over the last 50ns, is not sufficient to detect significant modification in the filament behavior. A reason is that the simulations have not reached equilibrium within the 200 ns, making average values non-representative. In addition, the NPFs are not homogeneous along their length since only a fraction of the filament proteins is bound to the D-loop; the rest interacts with the ssDNA or with the junction regions where the D-loop meets the relaxed dsDNA in B-form and forms a kink. This makes a global descriptor poorly adapted. We therefore turned to descriptors that can address the local level of monomer-monomer interfaces and capture trends in axis deviations. In recent work, Marien *et al*. introduced the Menger-based local LC and LF metrics and showed how that metrics encompasses the variability of both the curvature and the flexibility along protein chains, either folded or disordered [48], [49] (see **Supplementary information S1, Scheme S1C**). We reasoned that this metrics, when applied to the curvilinear axis associated to the protein filament, may inform about tension-associated trends occurring during the non-equilibrated simulation: either curvature collectively building or fluctuations unfolding within a relaxed state. **Figure 2** reports the evolution of the LF values along the NPF lengths for increasing D-loop lengths. The data indicates that the filaments conserve a strong rigidity up to 21bp, but start to fluctuate or become floppy as soon as the length exceeds 27bp. Interestingly, flexible regions in longer NPFs are mostly found in the D-loop binding filament turn that is closest to the 3’ extremity (interface numbers above 8 in **Figure 2A**). We also note that rigidity in the shortest filaments does not preclude curvature: for example in the 21bp simulation, first replica, curvature appears to build during the simulation, concentrated at particular filament levels (green to yellow shades in **Figure 2**, bottom panel). Time evolution of LC values for all D-loop lengths as well as LF data for the replica simulation can be found in Supplementary information **Figure S5**.

**Figure 2:**
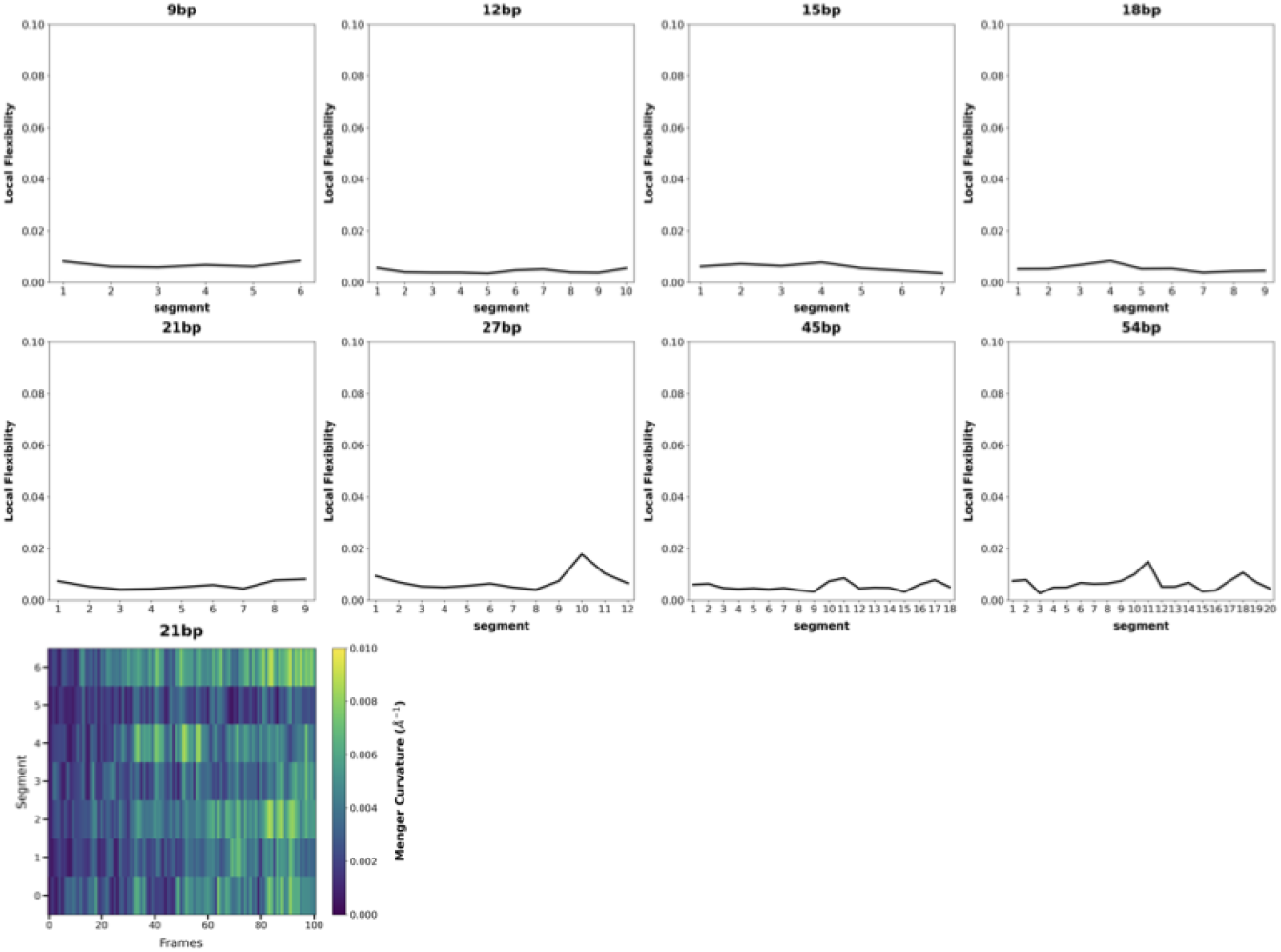
Fluctuations in the protein filament structures. (top) Time averaged LF values are reported in function of the position in the sequence of assembled proteins in the filament, with each number representing an interface between neighbor proteins. The numbers increase from the first interface in 3 ‘to the last interface in 5’. The results are given for several D-loop lengths between 9 and 54 bp. The bottom panel represents the time evolution of the local curvature LC value calculated for each interface of the 21bp model, during 200ns MD simulation (frames are analyzed every 2 ns); the color scale is displayed at the right of the plot.

To further characterize the filament overall distortions, we appealed to the elastic rod framework that can capture the filament deviations from a regular assembly in terms of bending, twisting or local curvature of the filament axis [47], [61]. Elastic rods axes were constructed from successive reference frames associated to pairs of consecutive subunits (**Scheme S1C**) and were regularly generated along molecular dynamics trajectories. **Figure S5** shows the time evolution of the parameters in function of the position along the filament (y-axis, numbered from 5’ to 3’, see schemes in **Figure S5** right panels) for three simulations (54bp, 18bp, 15bp, 1st replica). While, as already noted, the present simulations are too short to expect a full characterization of the NPFs internal mechanics, the results displayed in **Figure S5** demonstrate a strong potential for the elastic rod analysis to precisely dissect filament response in terms of curvature and twist, and detect filament regions that concentrates distortions.

### 3.3 Increasing the length of incorporated DNA modifies the protein-DNA interaction network

As already mentioned the models submitted to MD simulations initially present uniform geometries and similar networks of protein-DNA interactions in the filament region that binds the D-loop. **Figure 3A and 3B** respectively show the evolution of the distances between site I binding residues and the ssDNA phosphates, site II binding residues and the displaced strand phosphates, along the trajectory. We observe that the ssDNA backbone remains stably bound to site I for all simulated complexes whatever the D-loop length. To the contrary, **Figure 3C** shows a clear destabilizing effect of the D-loop length on the displaced strand binding to site II for D-loop lengths greater than 21bp, an effect which is remarkably reproduced in the simulation replicas (**Figures 3C and S7**.

**Figure 3:**
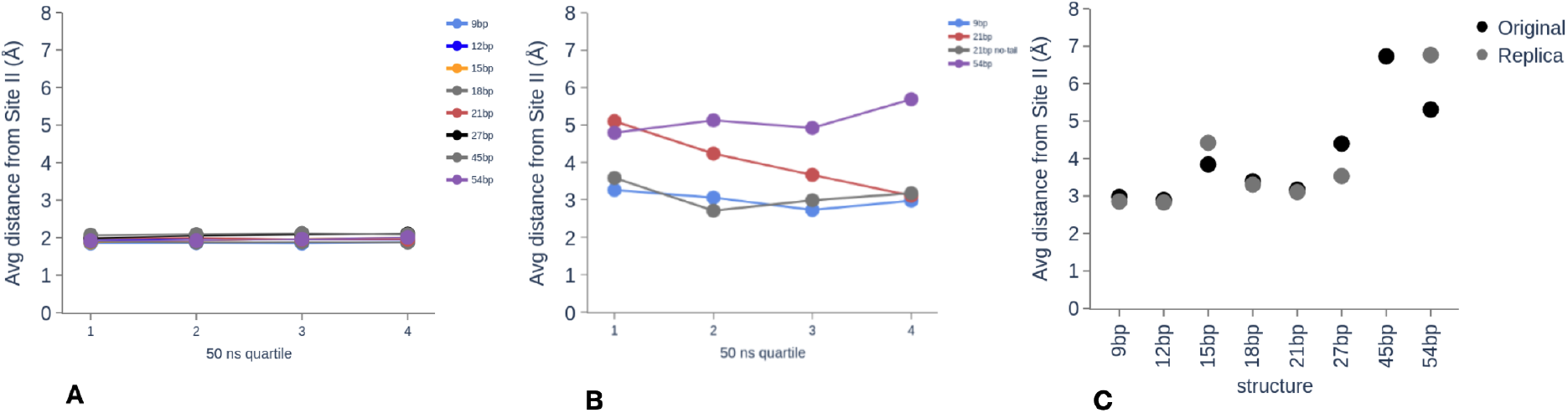
Time evolution of the average distances between residues defining the protein binding sites and phosphate groups from (A)the ssDNA bound to site I and (B) the displaced strand bound to site II. The distances are averaged over the DNA bases that belong to the D-loop and over successive 50-ns window quartiles of the 200 ns MD simulations systems with different D-loop lengths. Only the values concerning the first replica are displayed. In (A), the values are very close so that only two colors appear on the graph, other colors are hidden; in (B) the values concern three characteristic lengths of incorporated DNA: 9bp, 21bp and 54bp; two simulation results are displayed for D-loop lengths of 21bp, related either to the complete system (red) or a system when the B-form extremities of the dsDNA have been removed (grey); (C) DNA-length dependency of the average distance between residues defining the protein binding sites and phosphate groups from the displaced strand bound to site II, in the last 50-ns of the MD simulations; for each D-loop length, values for both replica are displayed, with different shades of grey. Site I and site II are respectively defined by residues 156-164, 210-212 for site I, residues 226, 227, 243, 245 for site II.

Below 21bp, we observe a small destabilization peak that culminates for 15bp, present for both replicas, suggesting that the 15bp model cannot fully conserve its outgoing strand contacts with site II. In contrast, the 21bp model reproducibly shows good adaptation. We will come back to that small peak in section 3.6. Looking now at the time response, we observe for the two independent simulations of the 21bp model that the contacts with site II are lost at the beginning of the production but get recovered during the course of the 200 ns trajectory (**Figure 3C** and supplementary **Figure S7**; the effect is more important for the first replica). Using extra care when restraining the outgoing strand DNA backbone during the model construction and minimization did not change that initial unbinding response (not shown). This observation suggests that for that specific length, accommodation of the outgoing strand binding in site II is obtained at the expense of a collective distortion of the filament structure. Alternatively, when we removed the dsDNA B-form extremities from the 21bp model, we observed that the displaced strand remained firmly bound to site II during the whole course of the simulation (**Figure 3B**, grey curve). This suggests that the presence of the dsDNA extremities anchored to the filament grooves in 5’ and 3’ of the D-loop is necessary for a full response of the filament.

We can now read **Figure 3C** in the perspective of the mechanical response of the NPF to building stress (**Figure 4**). If we exclude the 15bp system (grey rectangle), the curve can be interpreted as the transition from an elastic response below 27bp, where the system collectively responds to increasing stress to maintain its internal integrity, to a different state where the strong interactions between site II and the outgoing strand phosphates get broken, enabling the filament structure to relax. This transition can be related to the elastic to plastic transition that occurs in any material submitted to increasing stress, where internal contacts irreversibly reorganize in order to release the stress that can no more be handled by elastic deformation. However in our case, increasing the stress requires changing the system by increasing the D-loop length. We do not know in the present state whether a putative retro-transition from a 27bp D-loop to a 21bp D-loop, via a 24bp D-loop, would allow recovering the lost interactions in site II. Nevertheless, based on this analogy, we call the state with conserved contacts E-state and the state with reorganized protein-outgoing strand contacts P-state.

**Figure 4:**
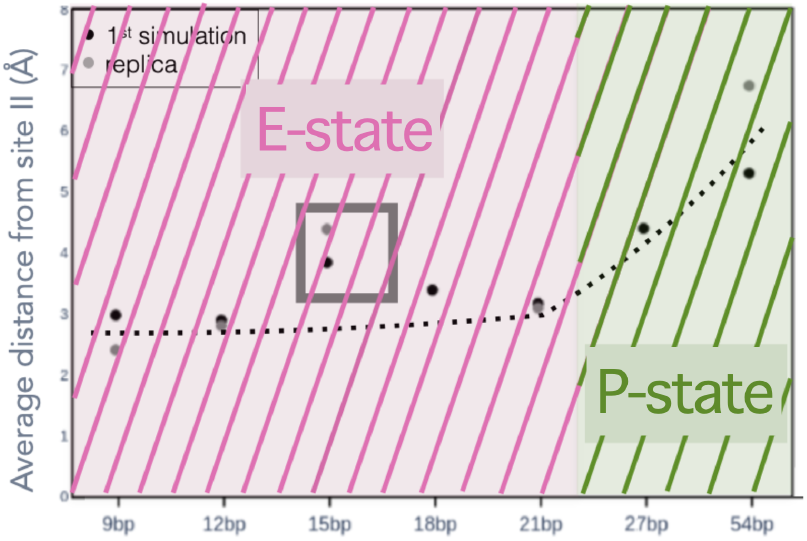
Response of the RecA nucleoprotein filament to increasing D-loop length. With the exception of the 15 bp model, the filaments with D-loop length lower than 27 bp (E-state) maintain their integrity in terms of internal contacts. For D-loop values higher than 27bp, the outgoing strand looses its connections with site II.

The numerical simulations offer privileged access to the details of that transition. For the shortest 9 and 12 bp models, phosphates from the displaced strand only punctually stray apart from their binding site II for durations that do not exceed 100 ns (**Figure S8**). Upon increasing the D-loop length, separations occur over longer time periods; finally beyond 27bp, contact losses spread over an increasing number of sites, to never be recovered over large regions of the filament, even though the formation and breaking of contacts remain dynamic. For all lengths, phosphates at both the 5’ and the 3’ extremities of the D-loop remain firmly bound to site II clusters of basic residues (dark blue regions in **Figure S8**).

### 3.4 From site II to site III

We wondered what happens to regions of the displaced strand that unbind from site II. To answer that question, we generated contact maps between each base of the displaced strand and protein residues in successive monomers. Maps for three representative models – 12bp, 21bp and 54 bp – are shown in **Figure 5**. Only the protein residues that contact the phosphate groups over more than 10% of the simulation time are represented; contacts appear as colored squares whose intensities depend on the fraction of the simulation where the contact is present. In these maps, we highlighted protein regions based on their close proximity to L2 loops, N-or C-terminal domains or site II, by shading them and lining the involved residues with different colors (see **Figure S9** for a 3D split-representation of those regions). Because both site II and the L2 loops define the displaced strand binding cleft, contacts with L2 loop residues situated in the (197-207) window are present even when the displaced strand is firmly bound to site II, as can be seen in the 12bp map.

**Figure 5:**
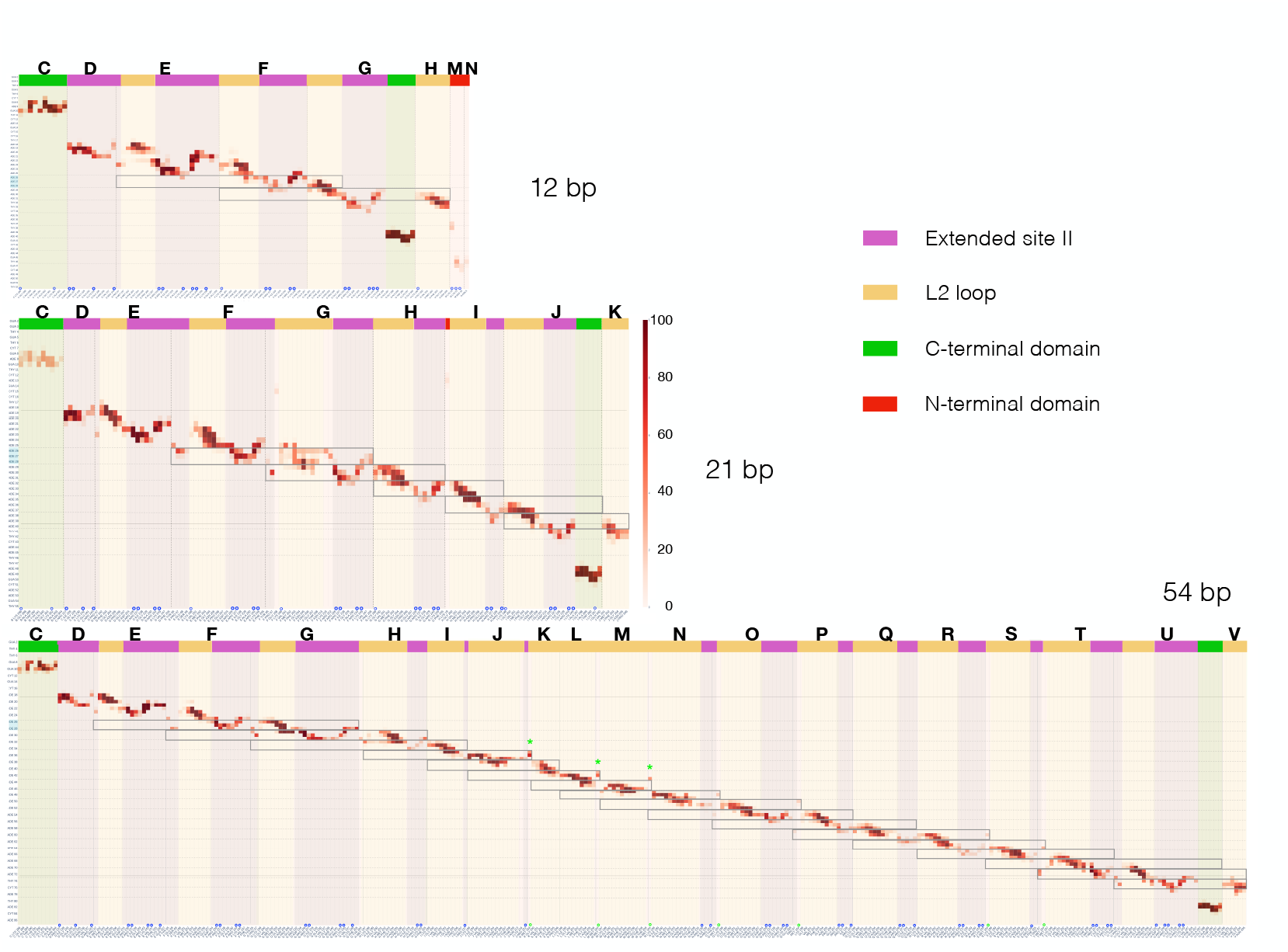
Contact map between phosphates in the displaced strand (y-axis) and protein residues (x-axis) from successive monomers in the 5’ to 3’ direction, averaged over 200ns MD simulations for three models with increasing D-loop size. Residues from different protein monomers are separated by broken lines. Different binding regions are delimited based on the residues belonging to or close to known domains (C- and N-terminal domain) or binding sites (L2 loop, extended site II), as defined in **Figure S9**.The residue names can be read by zooming on the figure in the numerical version. Blue circles mark the basic residues that bind DNA phosphate groups, green circles and green stars indicate contacts with methionine 164 which is part of the L1 loop in binding site I. Horizontal rectangles refer to the groups of three bases that were replicated from the bases numbered 26 to 28 in the 9bp model (directly issued from the CryoEM structure 7JY9) to construct models with increasing DNA lengths. We note that in the contact map of the 12bp, for which the filament extends in 3’ of the D-loop, the N-terminal domains of two monomers situated one helical turn after the 3’DBP contact the dsDNA 3’ extremity.

The map shows that contacts in the site II region extend over the whole four-strand *β*-sheet that contains the S2 hairpin and the site II basic residues Arg226, Arg227, Lys232, Arg243, Lys245 (referred to as “Extended site II” in **Figure S9**). While alternation between site II and L2 loop regions is regular or quasi-regular for the 12bp and 21bp models, we observe in the 54bp map that the extended site II region is either missing, or reduced to one weak residue contact, over a continuous region of the filament involving up to six consecutive monomers (one filament turn, corresponding to one third of the D-loop bound filament region). The same behavior, however with a shorter disconnected region, was observed when analyzing a former simulation of a 36bp D-loop model within a NPF binding the DinB polymerase at its 3’ extremity [37] (**Figure S10**).

In such region, some new L2 loop contacts are found, involving residues Thr208, Thr209, Asn213 and Lys216 that are generally not forming contacts when site II binding is strong. Also noticeable are the contacts with methionine 164 that appear in the regions where contacts with site II are lost. Methionine 164 belongs to the L1 loop and intercalates between site I-bound heteroduplex bases every three base pairs. The formation of contacts with Met164 is a good indicator that the displaced strand has come close to site I. Interestingly, **Figure S11** indicates that in those regions where the displaced strand disconnects from site II, its position with respect to the L2 loop locally shows good convergence with the position we previously proposed to be occupied by the complementary strand in the synaptic filament prior to strand exchange, *i*.*e*. site III [23] (see **Figure S2**). Our observations therefore indicate that as the D-loop length exceeds 27 bp, substantial regions of the displaced strand switch from site II to site III.

### 3.5 3’-5’ asymmetry of the D-loop interaction network

The network of protein-DNA interactions in the NPF can be divided into three groups: interaction of the B-form dsDNA extremity in the filament groove in 5’, involving its anchoring to a RecA C-terminal domain; interactions of the stretched-unwound three strands of the D-loop with binding sites I to IV ; interaction of the B-form dsDNA 3’ extremity in the filament groove in 3’, via its anchoring to the C-terminal domain of another monomer. In what follows, we will refer to the anchoring monomers in 3’ and 5’ as the 3’DBP and the 5’DBP (for DNA-binding protein). The spatial separation between the 3’DBP and the 5’DBP increases as the D-loop length increases. Here, we are mostly interested in the contribution of the displaced strand within this interaction network. At the short range level each monomer interacting in the center of the D-loop binds close to six bases, and each group of three bases mostly binds two consecutive monomers (**Figure 5**). This comes notably from the fact, already mentioned, that the S2 and L2 motifs that define site II at the level of a given base triplet belong to two adjacent monomers. Concerning the anchoring monomers, the extent of interactions with the DNA displaced strand differs between the 5’ and the 3’ extremities. In 5’, the proteins that bind respectively the B-form dsDNA extremity, the kink that separates the B-form dsDNA extremity from the D-loop, and the first DNA bases in the D-loop are distinct proteins (resp. *C, D* and *E* for all models in **Figure 6**, where *C* is the 5’DBP). These proteins also bridge the dsDNA 5’-extremity with the ssDNA region in 5’ of the the D-loop (**Figure S11**), however they remain disconnected from the D-loop region.

**Figure 6:**
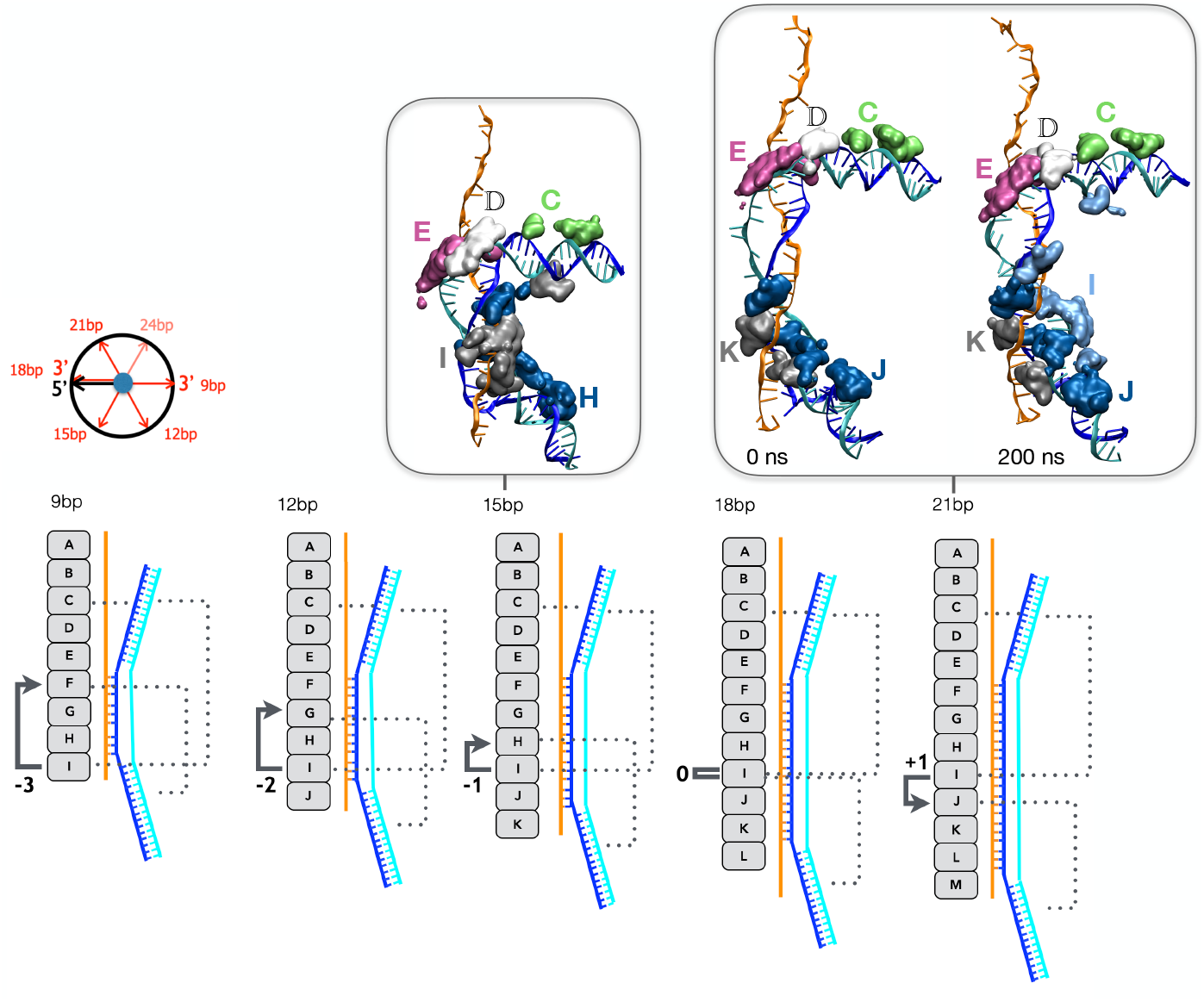
Scheme of the length-dependent evolution of the contact network involving RecA monomers, B-form dsDNA extremities and D-loop DNA strands for short D-loop lengths (< 21 incorporated base pairs). The scheme in the lower panel represents the sequence of RecA monomers (5’ to 3’ direction from top to bottom), for increasing lengths of the D-loop. Putative contacts with the B-form dsDNA extremities are shown with broken lines. Relative positions of the monomers that respectively bind the 5’ and the 3’ dsDNA extremities are marked with black arrows and their distance in terms of number of monomers is indicated. The scheme in the top left panel shows the relative directions of the 5’- and 3’-dsDNA extremities observed perpendicularly to the filament axis. 3D-snapshots from the MD simulations are displayed for the 15bp and the 21bp models; the DNA strands are in cartoon representation (same color code as in Figure 1) and protein regions that are closer than 7 Å from the dsDNA strands are shown in surface representation and colored according to the monomer they belong to.

In contrast in 3’, a same protein, the 3’DBP, simultaneously binds the B-form dsDNA extremity with its C-terminal domain, the kink region with its S2 hairpin and the second-last displaced base triplet with its L2 loop, which also intercalates between the heteroduplex second-last and third-last triplets; finally, the 3’DBP also binds the fourth-last heteroduplex base triplet with its L1 loop. These contacts are represented in dark blue in the inserts of **Figure 6** for both the 15bp and 21bp models, where the 3’-DBP corresponds to *H* and *J*, respectively. The 3’DBP shifts along the monomer sequence as more DNA triplets are incorporated in our model filaments, from monomer *F* in the 9bp model to monomer *U* in the 54bp model; however its contacts span up to three D-loop triplets regardless of the D-loop length, provided that that number of triplets is present in the D-loop in 3’ of the 3’-DBP: in the 12bp and the 9bp models, only two and one D-loop triplets, respectively, are available for binding. Finally, the monomer that binds the last triplet of the D-loop in 3’ immediately follows the 3’DBP in the 3’ direction. This provides intertwining within the protein-DNA contacts, which mostly involves the 3’ extremity.

### 3.6 Protein-DNA interaction network in the 15bp NPF

In addition to the interaction network described above, another type of contacts retained our attention.

A first type of contacts involves the monomer situated exactly six monomers in 3’ of the 5’DBP (monomer *I* in all models), and the B-form dsDNA extremity in 5’, that monomer *I* can potentially bind via basic residues of its N-terminal domain. In **Figure 6**, these basic residues appear as grey and light blue patches below the B-form dsDNA 5’ extremity in the 15bp and 21bp models, respectively. Such interaction involving the N-terminal domain of monomer *I* did not form immediately during the simulations but rather required some torsional adaptation of the simulated structures, as the N-terminal domain does not point exactly in the direction of the dsDNA 5’-extremity in the regular, initially built model. Nonetheless, that interaction was observed in all simulations (**Figure 7**), although in several cases it was mainly transient, sometimes appearing after several tenths of nanoseconds and persisting for time scales as small as nanoseconds (e.g. 54bp, **Figure 7**). In the transient cases, it may be considered as an opportunistic, weak interaction appearing according to the fluctuations of the C- and N-terminal domains. In some cases however, like the 9bp, 15bp (both replica) or 21bp (first replica) models, that interaction was much stronger and persisted over more than 100 ns. However the formation of contacts between the N-terminal domain of monomer *I* and the dsDNA 5’-extremity does not appear to be linked with the observed outgoing strand contact disruption since both the 15bp system (where central contacts of the outgoing strand are lost, **Figure S8**) and the 21bp system (where contacts are restored, **Figure 3B,C**) display strong 5’-DBP binding to the dsDNA extremity. **Figure 3B**.

**Figure 7:**
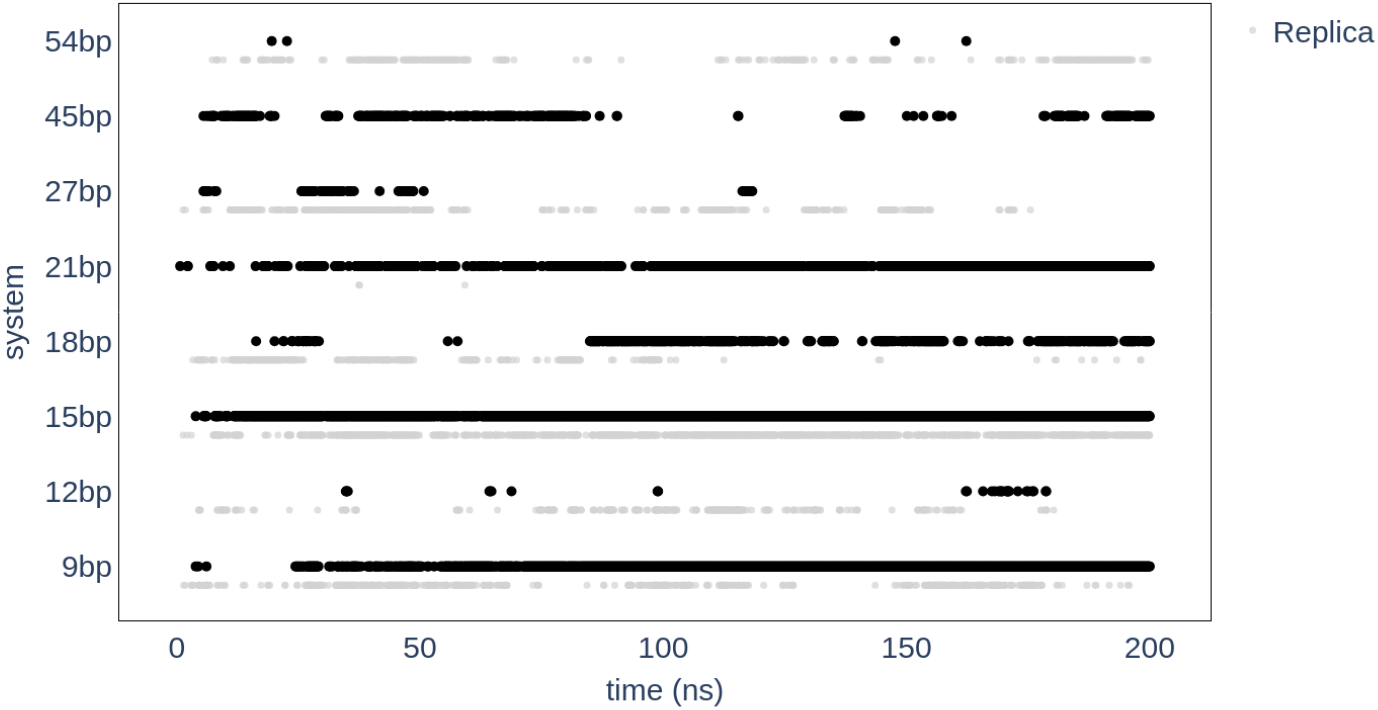
Time evolution of the interaction between the N-terminal domain of monomer I and the 5’-dsDNA B-form extremity for all simulations. A dot is drawn for each frame where the distance between two groups of atoms, namely residues 1 to 29 for I-N-terminal domain and the displaced strand bases 8 to 17 and Watson-Crick-bound bases in the complementary strand, drops below 5Å.

**Figure 6** illustrates another type of contacts that takes place at the opposite extremity of the D-loop. Quite surprisingly, for short D-loop sequences below 18bp we found that monomer *I* gets situated in 3’ of the 3’DBP monomer and separated from this monomer by up to 3 monomers that bind the D-loop (3, 2 and 1 separating monomers for the 9bp, 12bp and 15bp models, respectively). Such “retro-binding” may reinforce the already dense network of protein-DNA interactions and, when the interaction between monomer *I* and the 5’-dsDNA extremity is strong, produces a second level of intertwining interactions. Rigidification may ensue provided that the interaction with the D-loop is strong. We have noted that in the 9bp system, the 3’DBP protein only interacts with one triplet level of the D-loop; the number of spanned triplet levels, and therefore the strength of D-loop binding, increases with the D-loop size until it reaches its maximum value for 15bp and beyond. As the D-loop length increases above 15bp, monomer *I* gets positioned in 5’ of the 3’DBP with increasing separation, and the possibility of this second level of intertwining vanishes.

We now have a possible explanation for the curious observation of a secondary peak in **Figure 3B** for the two replica with 15bp D-loop length. We propose that in addition to the extensive interactions of the 3’DBP along three triplets of the D-loop, the reinforced interaction network due to retro-binding stiffens the filament so that it is no more compatible with a full binding of the displaced strand in site II. Conversely for the 21bp structure, the 3’DBP is now situated in 3’ of the 5’DBP, which releases some of the filament stiffness and makes the adaptation to site II binding possible.

Interaction of the 3’-dsDNA extremity with N-terminal domains of proteins across the groove is also possible as informed by the 12bp simulation. As mentioned in the Methods section, we built the 12bp model differently from the others by adding one filament turn in 3’ of the D-loop region and by lengthening the bound ssDNA accordingly; the model was otherwise constructed in exactly the same way as the other models. Given that the dsDNA can be incorporated anywhere within the recombination filament, the dsDNA 3’ extremity can insert in the filament groove during elongation as long as it does not reach the 3’ extremity of the ssDNA. In the CryoEM structure, the dsDNA 3’ extremity is bound at the filament 3’-end, which precludes observation of its interaction with the filament groove. We wanted to examine whether embedding the dsDNA 3’-extremity in a filament groove could influence the network of protein-DNA interactions within the filament. Observation of the contact map for the 12bp model in **Figure 4**, top, provides an element of response, as contacts can be detected between the displaced strand and the N-terminal basic residues Lys19, Lys23 and Arg33 of monomer *M*, situated exactly 6 monomers after the 3’-dsDNA-binding monomer *G* in the 3’ direction. Monomer *N* is also involved with a very weak contact of residue Asn5 of its N-terminal domain. These contacts only involve the 3’-dsDNA extremity and not the D-loop, so they probably do not take part in the effect of the D-loop length on the filament properties. However, they may be involved in strand exchange progression.

### 3.7 Increasing the length of incorporated DNA only slightly modifies the protein-protein interaction network

Protein-protein interfaces naturally fluctuate due to thermal motion, which results in the residue contact network across the interfaces undergoing dynamic exchanges. It was shown that for stable complexes, such exchanges does not involve more than half the initial contacts, or in other words, the f_NAT_ value, where f_NAT_ is the ratio of conserved contacts, does not decrease below 0.5 [60]. This observation also holds for protein-protein interfaces that are embedded in a filament, as discussed in [62]. For protein filaments under tension, thermal fluctuations may be modulated by the filament response, which may concentrate the load on the weakest points of the filament structure, the interfaces. Based on the observations described in the previous sections, concerning the 5’-3’ asymmetry in the protein-DNA contacts, we could have expected that the load would be distributed unevenly among interfaces between monomers that differently bind the D-loop and the B-form dsDNA anchors.

**Figure 8** shows the f_NAT_ values averaged over the last 50 ns of the MD simulations for each interface of the models with varying lengths. Most interfaces remained pretty stable during the simulations. We observed however one occurrence of an interface shift with f_NAT_ values below 0.4, for one replica of the 18bp model and for the interface between proteins that bind the 5’-B-form dsDNA extremity (see **Figure S13**). For the other models, we note a trend towards lower f_*NAT*_ values for monomer interfaces close to the 5’-kink. However, the effect is too weak for us to draw a conclusion, which could be expected given the short simulation time: disrupting stable contacts and forming new ones are energy and time-consuming events. Longer simulation times would be necessary to obtain meaningful information.

**Figure 8:**
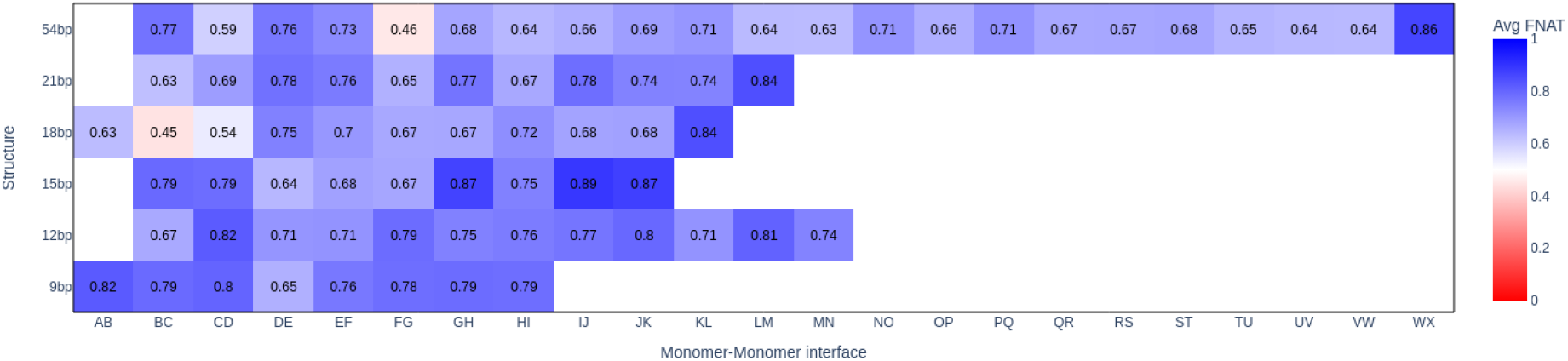
Heatmap of average interface f_NAT_ scores over the last 50 ns of 200ns simulation for each system. The color scale ranges from red (low f_NAT_) to blue (high f_NAT_). f_NAT_ value below 0.5 is typically interpreted as a significant disruption of the interface.

## 4 Discussion and conclusion

In this study, we used non-equilibrium molecular dynamic simulations to examine the response of the RecA nucleoprotein filament to increasing lengths of bound D-loop. The first noticeable information we came to is that the filament is indeed sensitive to the length of the D-loop it binds to and that it responds differently to increasing D-loop lengths. Broadly, our analyses show that up to a threshold length that we evaluate at 27 incorporated base pairs, the response is elastic and collective: the filament as a whole adapts to the added stress by increasing its overall deformations thus preserving the DNA binding interactions. After that threshold, the response involves significant rewiring of the protein-DNA interaction network inside the filament. Possible consequences for the homologous recombination process will be discussed below.

First we need to refine the broad-stroke picture described above by dissecting how different types of interaction define the nucleoprotein filament response. The present analyses, and notably **Figure 3A**, strongly suggest that the sensitivity to added DNA length can be quasi-exclusively attributed to the displaced strand and its binding to the secondary DNA binding site. As already mentioned, the clusters of basic residues defining site II are laterally displaced from the axis and are separated from binding site I by a continuous array of stacked L2 loops from successive protein monomers [14], [63]. That topology imposed by the NPF induces an extreme stress in the displaced strand backbone. Our results indicate that the sensitivity to that stress requires that the dsDNA B-form extremities are anchored in the filament groove.

The heteroduplex bound to site I has a strong stiffening effect on the filament. As discussed by Hegner and coll. [54] who characterized it experimentally, that stiffening effect cannot be attributed to either the protein or DNA components, but rather to their binding. Yet, our present simulations show that increasing the size of the heteroduplex in site I does not modify the connectivity network in site I, which remains highly regular for all models. On the DNA point of view, this observation is in line with early characterization of the site I-bound heteroduplex structure as a metastable state in the structural transition of double-stranded DNA under stretching load [64], [65]. Regular intercalation of hydrophobic side chains every three base pairs stabilizes that metastable state and indeed, our previous work suggested that once it is formed, the bound heteroduplex can retain its structure even in the presence of mismatches [28]. As former observations also pointed out, the two side chains that intercalate at each inter triplet site are provided by two different (successive) monomers, which contributes to knitting a dense network of local interactions. We can securely infer that that dense network, while it certainly contributes to the overall filament stiffening, does not allow forces to be transmitted over more that 3 or 6 base pairs in site I, in line with former single molecule observations about the exceptional stability of RecA-bound dsDNA and its insensitivity to changes in tension [30]. To the contrary the binding of the outgoing strand in site II relies on a set of successive lateral contacts. Although those contacts are individually very strong, they can break when submitted to load that exceeds a given value. In an analogy with the macroscopic world, the contact network in site I may be compared to a zip fastener while the contacts in site II would be an alignment of snap buttons. Once the load gets too high, the snap buttons can collectively break open.

We observed what could be compared to such a break opening for D-loop lengths greater than 27bp. While rupture of site II contacts was found to be limited in time and space for smaller lengths, and could be assimilated to thermal moves, rupture events in longer D-loops appeared to spread along the sequence and their duration increased. Soon, regions of the displaced strand exceeding 10 consecutive bases unbound from site II and switched to site III. The location of those regions along the sequence remained submitted to variations, as a result of the filament global fluctuations; however full binding to site II was never recovered during our simulations for D-loop lengths greater than 27bp, while full recovery could be observed for the 21bp simulation in spite of the binding being incompatible with the straight regular filament characterizing the starting model. We also note that in all simulated cases, the displaced strand conserved strong interactions with site II contacts at both the 5’ and the 3’ extremities of the D-loop.

In the absence of experimental information on the strand arrangement within RecA-bound D-loops, we can only speculate how a partial switch to site III may interfere with the HR process. By loosening the displaced strand, it may permit the incorporation of more triplets in the D-loop without destabilizing the assembly; by occupying site III, the displaced strand may inhibit potential binding of the complementary strand that would result from reverse pairing exchange. This would make filament disassembly in case of reaching heterologous dsDNA regions impossible for longer filaments in the absence of ATP hydrolysis.

Our work further points to significant differences in the strand and monomers wiring between the 5’ and the 3’ extremities of the D-loop, which implies strong consequences in terms of collective behaviors at each extremity. It has been established that the geometry of the junction, or kink, between the B-form dsDNA extremity in 5’ and the stretched and unwound D-loop region differs from the geometry of the junction in 3’ [23], [29] (note that in reference [29], the kink in 5’ is called 3’-tilt and the kink in 3’ is called 5’-tilt). In 3’, the B-form dsDNA forms an angle of about 45° with the filament while the angle in 5’ amounts to about 90°, in agreement with early AFM observations of Rad51-DNA filaments [17]. This difference reflects a drastic change in properties when it comes to internal connections: with the exception of monomer *I* whose interaction via its N-terminal domain shows variable levels, the proteins that bind the B-form DNA extremity and the junction in 5’ do not interact with the D-loop; conversely, the proteins than bind the B-form DNA and the junction in 3’ participate to an intricate network of interactions that involves a D-loop region that extends more than half an helical turn. We can therefore expect the transmission of forces to be very different in 5’ and in 3’, with stronger collective behavior in 3’, while the 5’ dsDNA B-form extremity would be mostly uncorrelated from the rest of the dsDNA. Although of a different nature, asymmetry in the filament behavior has already been documented in a former exploratory modeling study where we assessed the effect of modifying one protein-protein interface as a result of ATP hydrolysis. In that study, topological reorganization of the filament groove occurred mostly in 3’ of the modified interface and could span one filament turn [62].

At this point, we have shown that the filament responds to the increasing stress in the displaced strand of the D-loop, and we have dissected how forces can be transmitted inside the filament. Although it is outside the scope of the present study, the question of ATP hydrolysis and its consequences in terms of filament dissociation remains pending. In RecA filaments that do not undergo strand exchange, ATP hydrolysis has been shown to result in monomer unbinding only at the filament extremities, with a probability much higher at the 5’ extremity. However, our former work showed that when strand exchange is in progress, ATP hydrolysis results in the DNA completely dissociating from the NPF in 5’ of the D-loop, therefore at the heart of the filament, for D-loop length as short as 20 base pairs [66]. Such feature was observed only in filaments that undergo strand exchange [67]. Our simulations using classical force fields did not mean to address ATP hydrolysis and were clearly much too short for large disturbance to be observed in the protein-protein interfaces. However, observations from the present work lead us to emit some guesses. One such guess is that interface dissociation would be more prone to occur in the vicinity of the 5’ kink, where the connection with the D-loop is weak, than in the D-loop 3’-extremity that harbour dense connectivity network. Another guess is that such dissociation may be favored in shorter filaments where the tension in the filament is stronger, which may locally destabilize RecA-RecA interfaces. However, in the absence of specific experimental observation the present discussion can only be considered as mere speculation, which merits being pursued in separate studies.

Finally, the observations reported in this study may help understand mechanosensible processes that take place within other collective protein filaments [7]. Indeed, the nucleoprotein filament of homologous recombination offers a unique example of a macromolecular system that spontaneously builds internal stress by successive addition of discrete load quantities. The fact that we could access its mechanical properties through numerical simulations at an affordable computational cost opens new perspective for assessing the role of protein filaments in force sensing. The tools and analyses we have developed for this study may prove useful in this regard.

**Notes** All trajectories have been deposited on Zenodo (doi:10.5281/zenodo.18377475)

## Author contributions

Original draft preparation, A.S. and C.P.; Review and editing, A.S., M.P., C.D, S.N, A.D. and C.P.; Visualization, A.S., M.P., A.D. and C.P; Supervision, C.P. All authors have read and agreed to the published version of the manuscript.

