## Supplementary Figures S1 to S11 for "Mechanical response of the RecA nucleoprotein filament to increasing D-loop length"

### Mechanosensitivity of the RecA nucleoprotein filaments of recombination: Response to increasing the length of incorporated DNA

Afra Sabei, Alex Détruit, Sébastien Neukirch, Claudia Danilowicz, Mara Prentiss and Chantal Prévost

**Supplementary information SI1** - Definition of axis segments and axis frame; Schemes S1A and S1B

**Figure S2** - DNA binding sites in the RecA nucleoprotein filaments of recombination

**Figure S3** - Mapping the protein-protein and protein-DNA interactions within D-loop-bound RecA nucleoprotein filament

**Figure S4** - Time response of the axis shortening descriptor.

**Figure S5** - Time evolution of the Local Menger curvature during 200 ns MD simulation

**Figure S6** - Evolution of the elastic rod parameters

**Figure S7** - Time evolution of the distance between site II and the displaced strand phosphate groups

**Figure S8** - Sequence dependent evolution of the distance from site II of the displaced strand phosphate groups

**Figure S9** - DNA binding motifs in the RecA nucleoprotein filaments of recombination

**Figure S10** - Contact maps for the 15bp and 36bp models

**Figure S11** - Site II versus site III occupancy during 200-ns MD simulation

**Figure S12** - D-loop binding regions of the filament monomers

**Figure S13** - Evolution of Fnat, the fraction of conserved interface contact pairs during 200ns of MD simulation of the 18bp model

### Supplementary information S1 - Axis frame definition

#### A. Broken lines: definition of segment axes

For each pair of interacting proteins:

1. Define the parameters of the screw transformation:axis **Ω** ; translation ***trans***
2. Define the centers of mass of each protein P1 and P2: $C_{M1}$ and $C_{M2}$
   $C_M$ middle point between $C_{M1}$, $C_{M2}$
3. Define M as the projection of $C_M$ on **Ω**
4. Define the axis segment related to the P1, P2 interface
   The axis segment is centered at M and its length is ***trans***

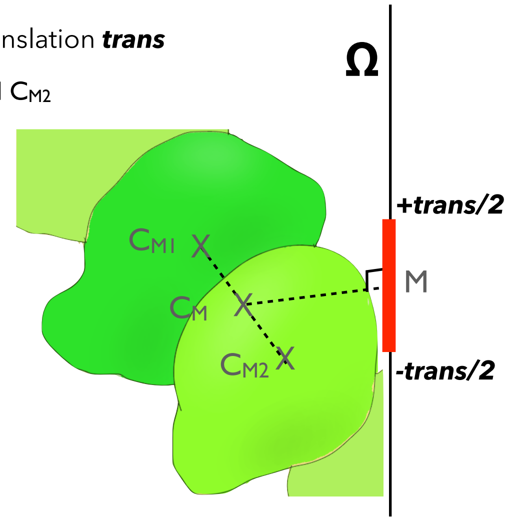


Scheme S1A

#### B. Elastic rods: definition of successive frames

For each pair of interacting proteins:

1. Define the parameters of the screw transformation: axis **Ω** ; unit vecteur **u**
2. Define the centers of mass of each protein: $C_{M1}$ and $C_{M2}$
   $C_M$ middle point between $C_{M1}$, $C_{M2}$
3. M projection de $C_M$ sur l'axe
   **u** = axis vector
   **v** = **M C**$_M$ / || **M C**$_M$||
   **w** = **u** X **v**

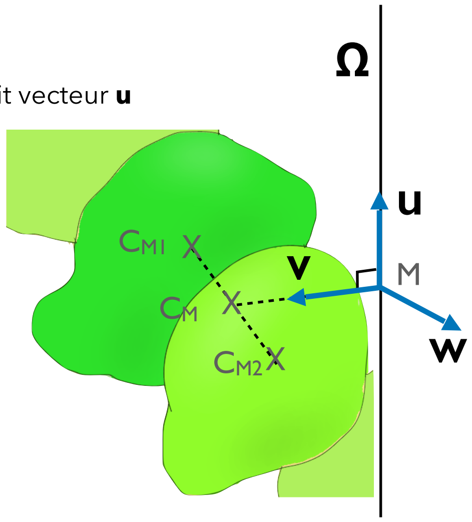


Scheme S1B

#### C. Menger curvature and flexibility

Given a curvilinear axis that links the M points defined above, local curvatures and flexibilities were defined as follows:

**PMC** = $1/R$, where **PMC** is the instantaneous local curvature

**LC** = <**PMC**>, where the average is taken on the whole trajectory and **LF** is the standard deviation.

A spacing value $i = 2$ was selected based on the comparison of **LC** and **LF** values obtained for $i = 1, 2$ or $3$

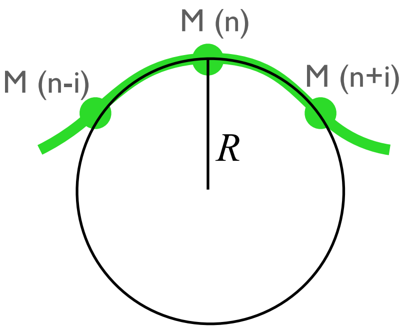


Scheme S1C

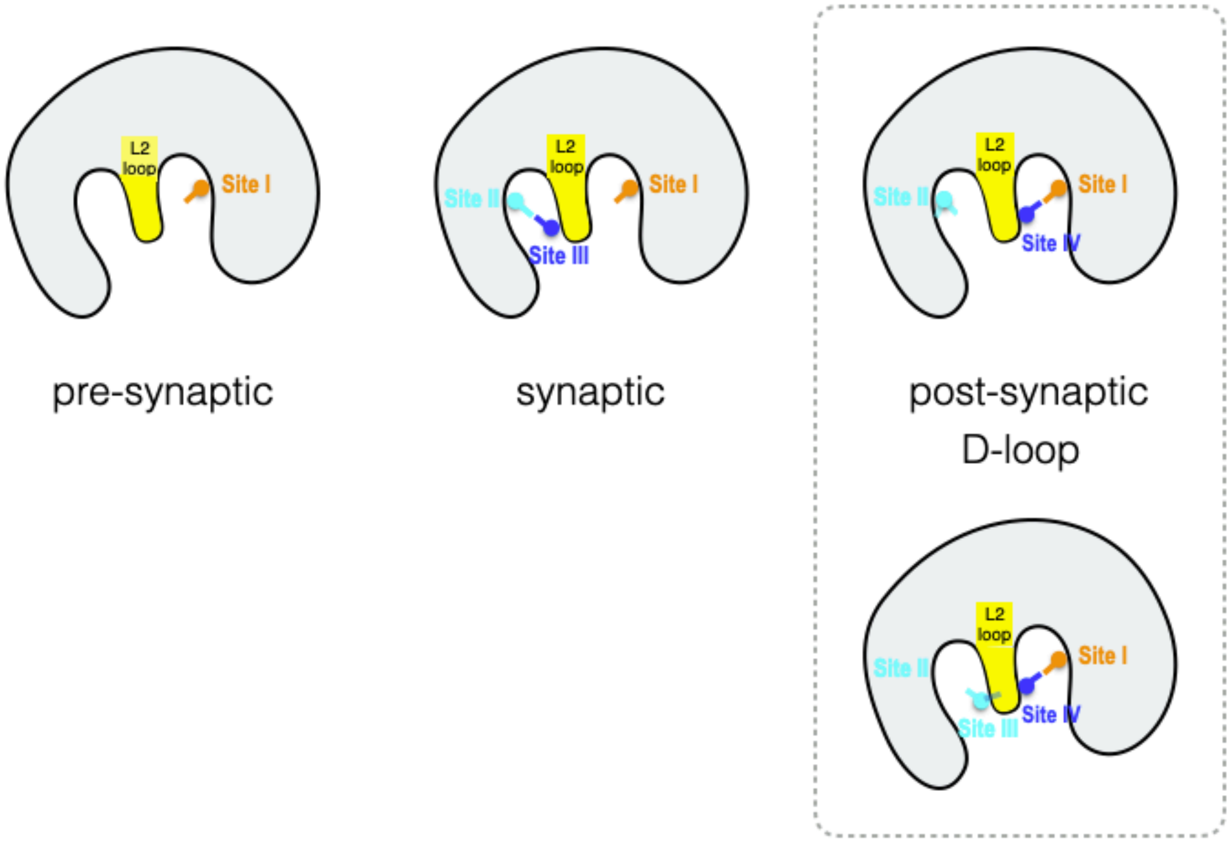


**Figure S2** - DNA binding sites in the RecA nucleoprotein filaments of recombination

The scheme represent slices of the RecA nucleoprotein filament taken perpendicularly to the filament axis. In the pre-synaptic filament, the incoming ssDNA is bound in site I; the synaptic filament is a transient intermediate where the searched dsDNA is bound to site II (outgoing strand) and III (complementary strand); in the post-synaptic filament, the complementary strand has relocated to site IV to bind the ssDNA in site I; the outgoing strand remains in site II in short D-loops but partly relocates to site III in longer D-loops (this work). The flexible and bulky L2 loop that separates site I and IV from site II and III has been colored yellow.

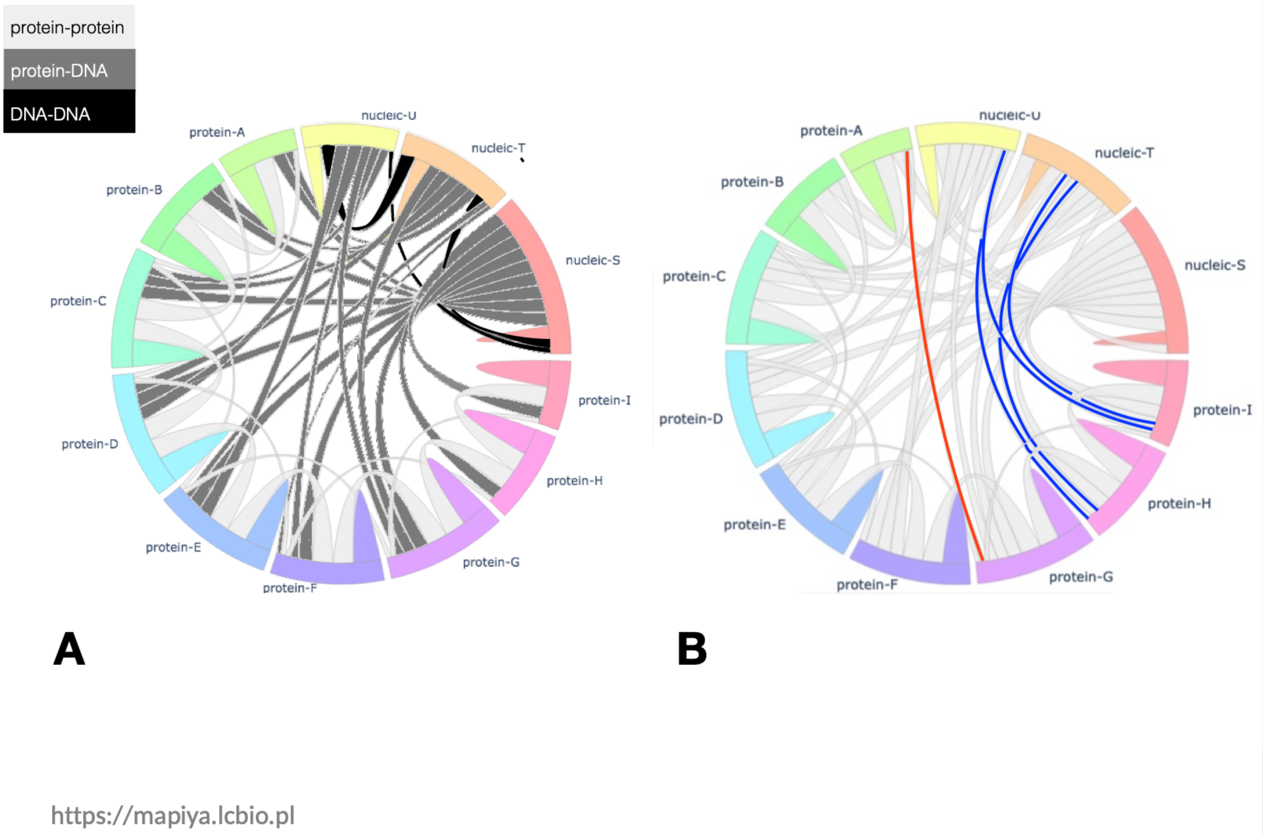


**Figure S3 -** Mapping the protein-protein and protein-DNA interactions within D-loop-bound RecA nucleoprotein filament

**Contact maps** between the RecA monomers and the DNA strands generated by the Mapiya web server (*https://mapiya.lcbio.pl*) [A. E. Badaczewska-Dawid, C. Nithin, K. Wroblewski, M. Kurcinski, and S. Kmiecik, Nucleic Acids Res, 2022 50(W1):W474-W482, DOI: 10.1093/nar/gkac307] for (A) the CryoEM structure, PDB code 7jy9 (B) the 9bp model derived from the CryoEM structure after 200 ns molecular dynamics simulation. Each ribbon in the map that links two molecules indicates contacts between these molecules, with the width of the ribbon increasing with the number of contacts. In (A), we have colored in white, grey and black the protein-protein, protein-DNA and DNA-DNA contacts. We note that although the D-loop only extends along 3 monomers D, E F (corresponding to 9 bp), interactions of the dsDNA B-form regions with C-terminal domains in the filament groove involves additional four monomers, three of them (A,B,C) in 5' and one (G) in 3' of the D-loop. In (B), the red ribbons indicate new protein-protein contacts that form during MD simulations between proteins situated across the filament groove; the blue ribbons feature new, transient protein-DNA contacts that involve the B-form DNA tail region non-specifically bound to protein C-terminal domains on 5’ (proteins B, C) and proteins H and I across the filament groove.

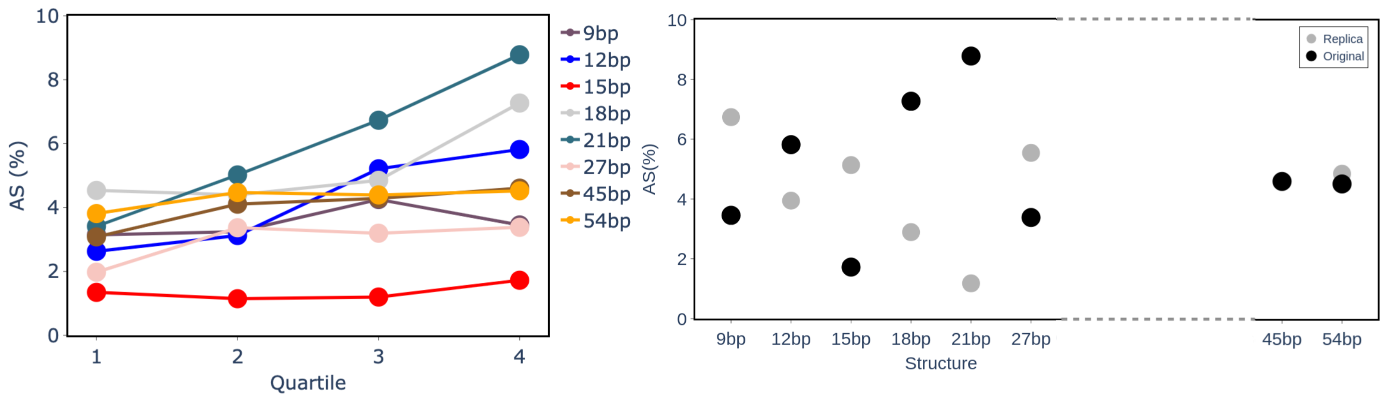


**Figure S4** - Time response of the axis shortening descriptor during 200 ns MD simulations. (Left) Evolution of the AS values averaged over successive 50-ns time windows (the quartiles); each color is associated with a different D-loop length, indicated at the upper right of the panel (right) AS values of the last quartile (average value over the last 50 ns) as a function of the D-loop length; black and grey circles represent values from two independent simulations, with the first one corresponding to values displayed the left panel. A value of 0 for *AS* indicates that the helix shows no distortion.

As could be expected, *AS* values depart from 0 due to thermal moves; however while they fluctuate for the 9bp, 45 or 54bp models (Figure S4A), *AS* values were found to continuously increase during the 200 ns of the 21bp first replica simulation, which may signal adaptation of the filament to strong internal constraints. However, this behavior was not reproduced in the replica 2 simulation of the 21bp model, although visual examination (not shown) also indicated some degree of collective curvature building. Lateral deviations between axes segments as well as variations in the segment lengths, together with the short out-of-equilibrium simulation time are probably responsible for the *AS* metrics poorly reflecting these trends: although some characteristics were retained in the two replica, such as the fluctuating values for 9bp and 54bp, the continuously increasing value for 21bp and the low value for the 27bp simulation, figure S4B shows no convergence of the *AS* values of short D-loops. For that reason, we turned to the Menger curvature and flexibility metrics that much better capture the visual behavior (see main manuscript and Figure S5 C).

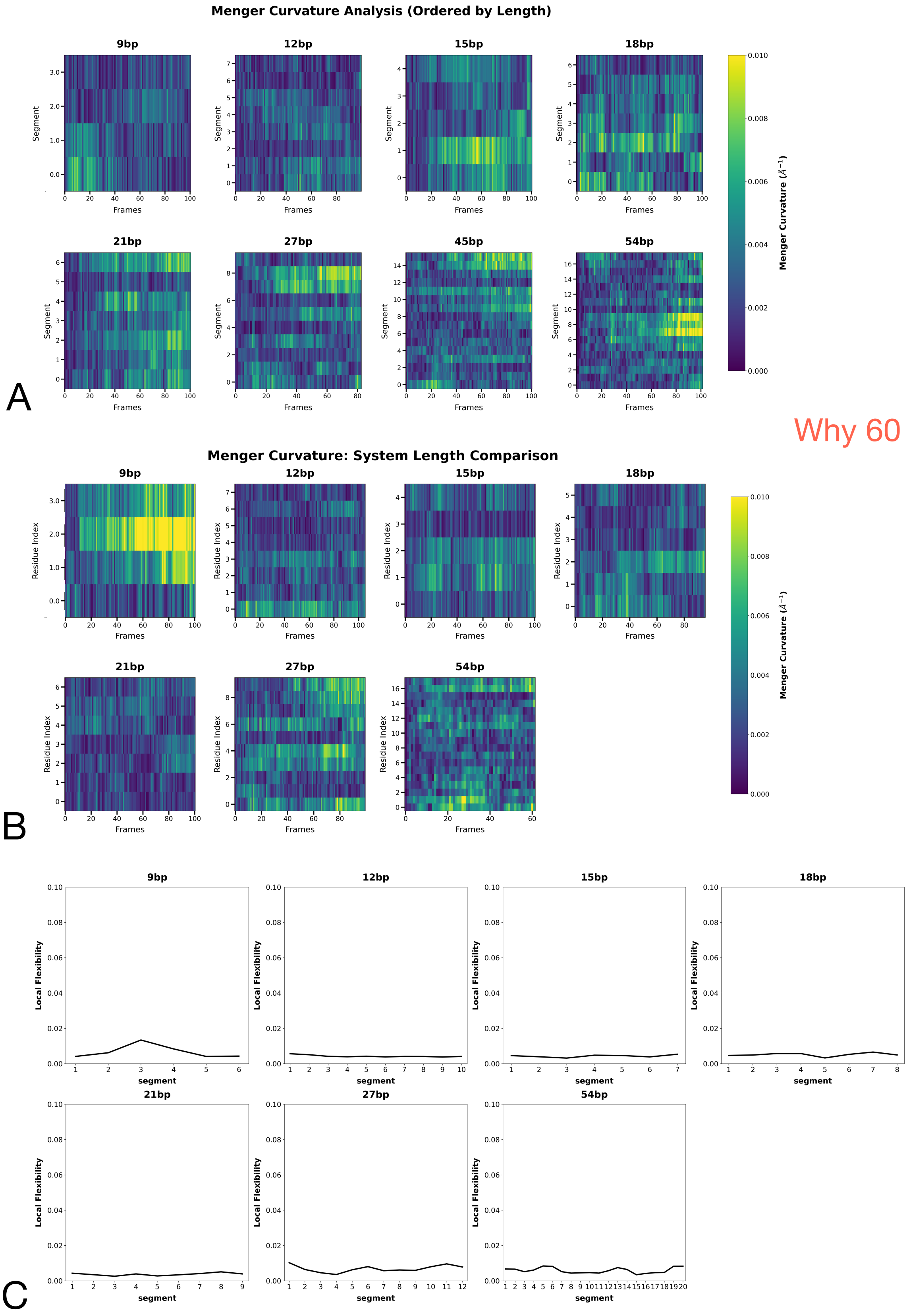


Why 60 frames?

**Figure S5 -** Time evolution of the Local Menger curvature LC (A, B) and the local flexibility LF (C) during 200 ns MD simulation for two series of simulation, replica 1 (A) and 2 (B, C); frames were analyzed every 20ns. Variation of LC values follows the color scale represented at the right side of each panel. The LC and LF values were calculated with the separation value *i* = 2 (see Scheme 1C), which means that the curvature is calculated over a five-point segment centered at the value corresponding to the residue index.

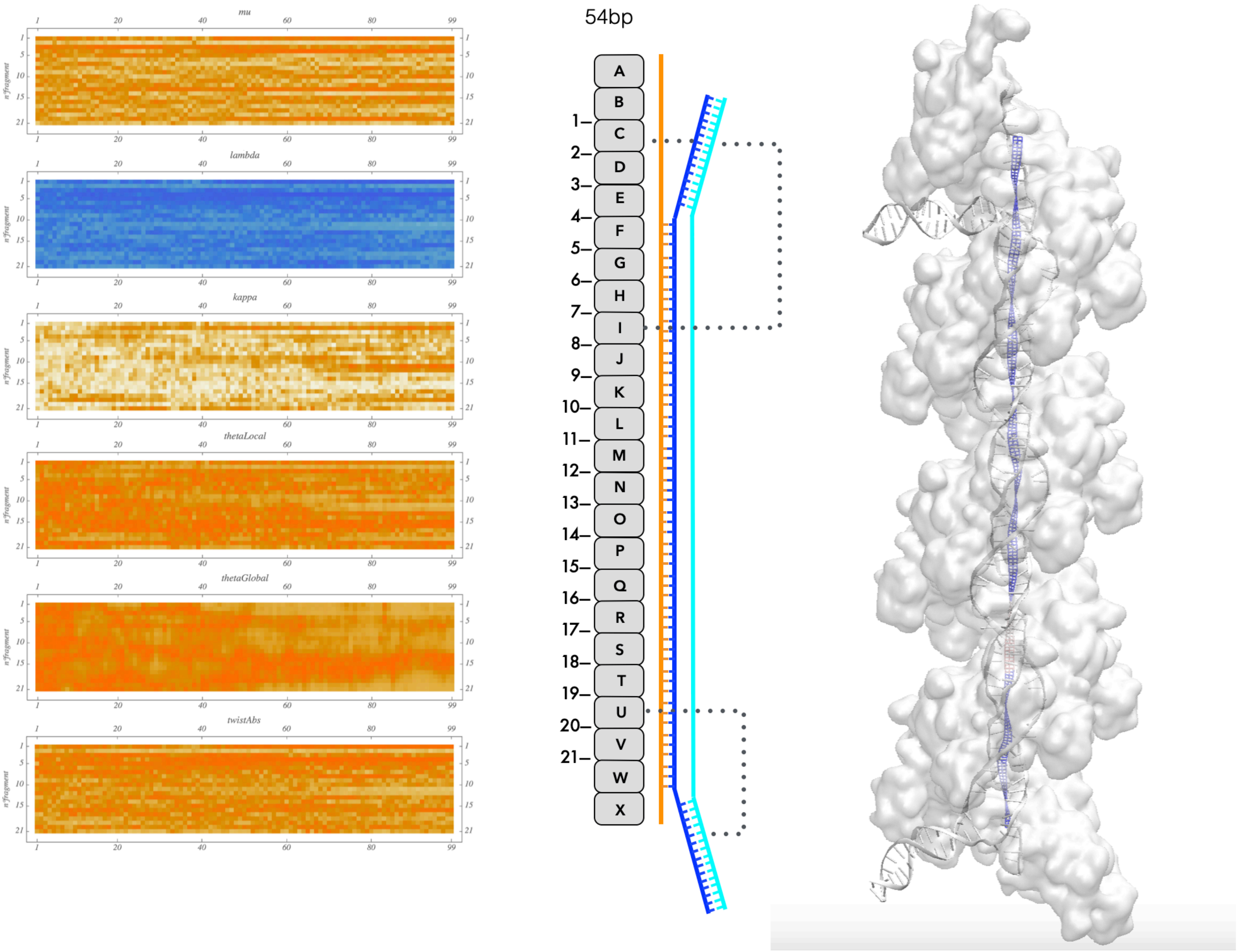


**A**

**Figure S6 - Evolution of the elastic rod parameters.** for (A) the 54bp model, (B) the 18bp model and (C ) the 15bp model. (left) evolution of the mechanical parameters describing the filament axis at each protein-protein interface level (y-axis) during 200ns MD simulation, replica 1; frames were analyzed every 20ns; description of the parameters can be found below; (right) schematic representation of spatial arrangement and the internal connectivities between the protein monomers and the three DNA strands (ssDNA in orange, bound to site I; complementary strand in blue; displaced strand in cyan, bound to site II in the D-loop region where the pairing has been exchanged); in (A), a structural view of a nucleoprotein filment snapshot (white, transparent) figuring the elastic rod axis in the center has been added for comparison; the rod representation was designed in order to visualize the torsional variations; color shading enable visualizing the variations of chosen parameters (here θGlobal; deviations increase from blue to red).

- $\mu = 1/\sqrt{T}$ : scaling factor, link to the norm T of the force vector.
- $\lambda$ : projection of the moment on the force axis
- $\kappa L$ : average curvature of the fragment.
- $\theta$Local : projection of the force vector on a local vector that characterizes the direction of the elastic rod fragment.
- $\theta$Global : projection of the force vector on a global vector that characterizes the direction of the structure.
- twistAbs : absolute torsion of the fragment.

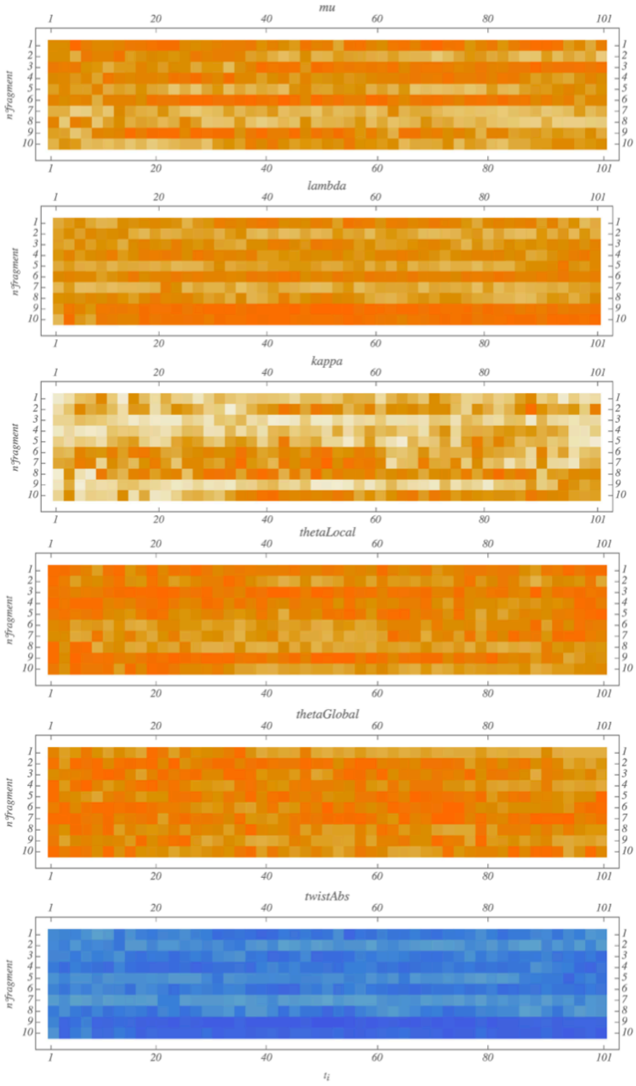


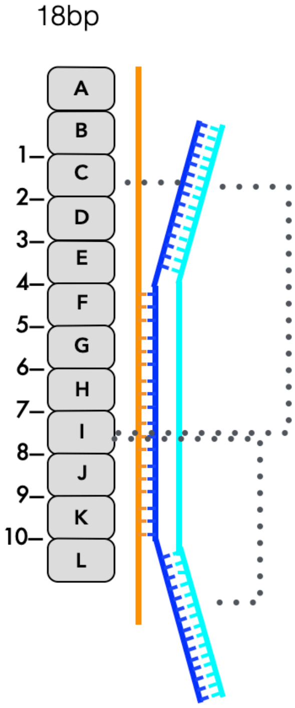


**B**

*(Figure S6 - continued)*

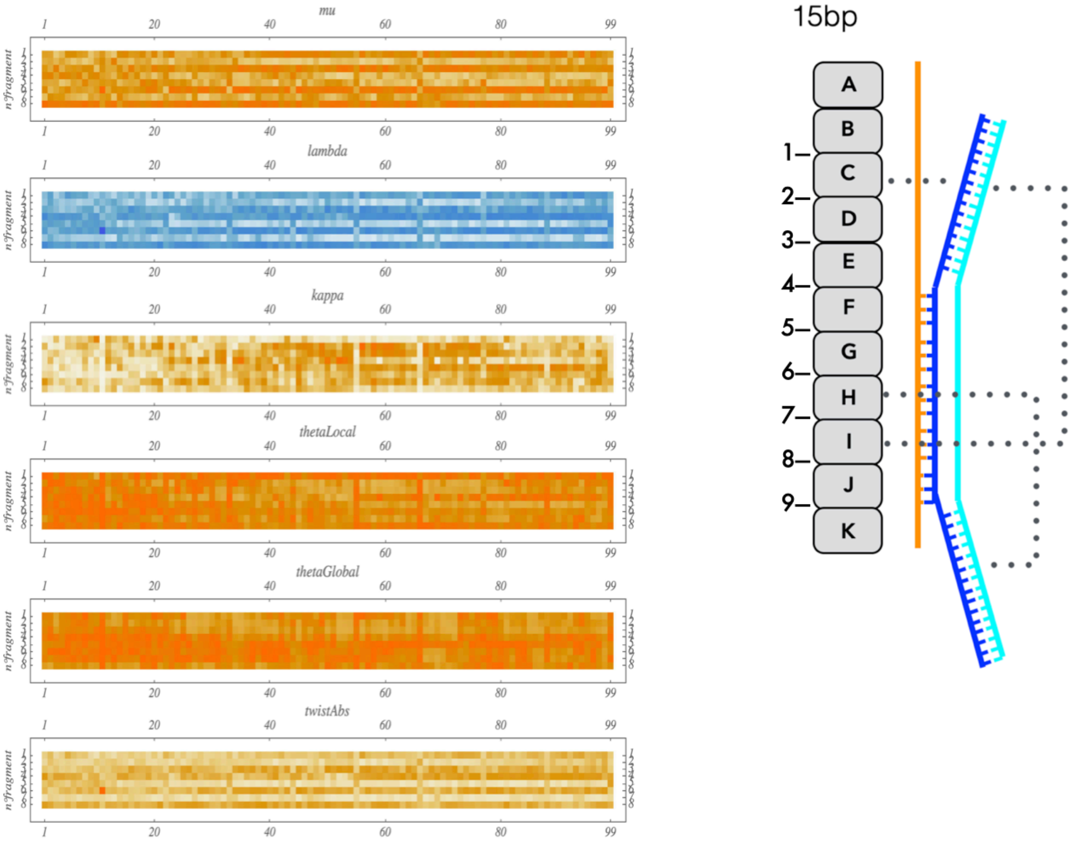


C

(*Figure S6 - continued*)

**Additional comments:** The first two parameters concern the evolution of the force and its moment, displayed along the filament. For the 15bp and the 18bp models, both force and moment are concentrated on specific interfaces, that are not necessarily identical (for example in the 15bp simulation, a high moment at interface 4 is not associated to a high force, which rather concentrates on interface 3). For the 54bp simulation, distribution of the force and moment along the filament is more diffuse and fluctuates. In all cases, the moment is almost completely transmitted to the Twist parameter in terms of localisation in the filament interface sequence, while curvature can  take place in regions that differ from the force localization. For the 15bp and the 18bp models, twist distortions enable interactions between the monomer I N-terminal domain and the dsDNA 5’ extremity to form (see Figure 7, main test).

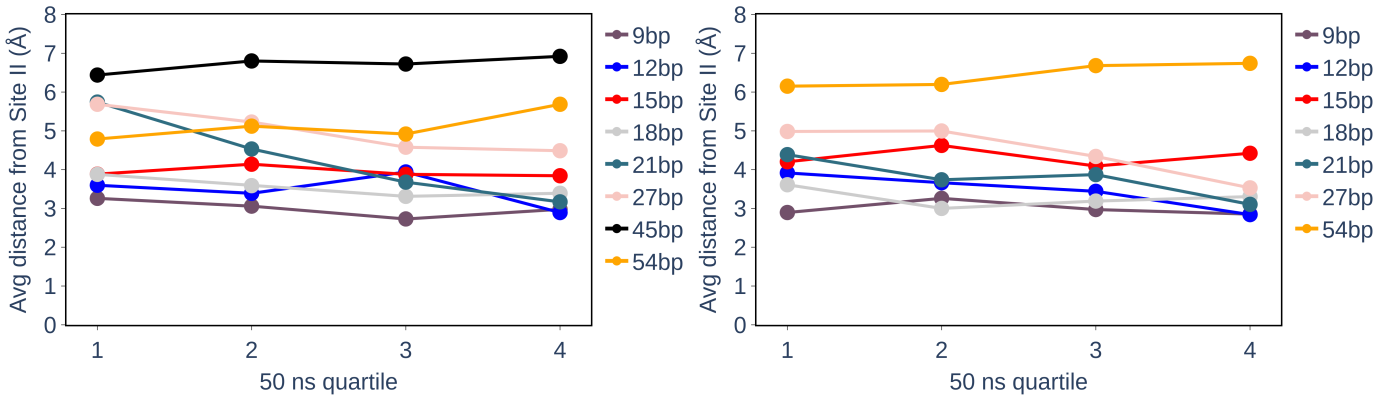


**Figure S7 -** Time evolution of the distance between site II and the displaced strand phosphate groups

Same as in Figure 3C. The left and right panels respectively represents the distance evolution for the two replica. For a given outgoing strand phosphate group belonging to the D-loop, the reported distance is taken as the shortest distance between that group and protein residues 226, 227, 243, 245. Each point represents the average distance over 50ns and over all outgoing strand phosphates belonging to the D-loop.

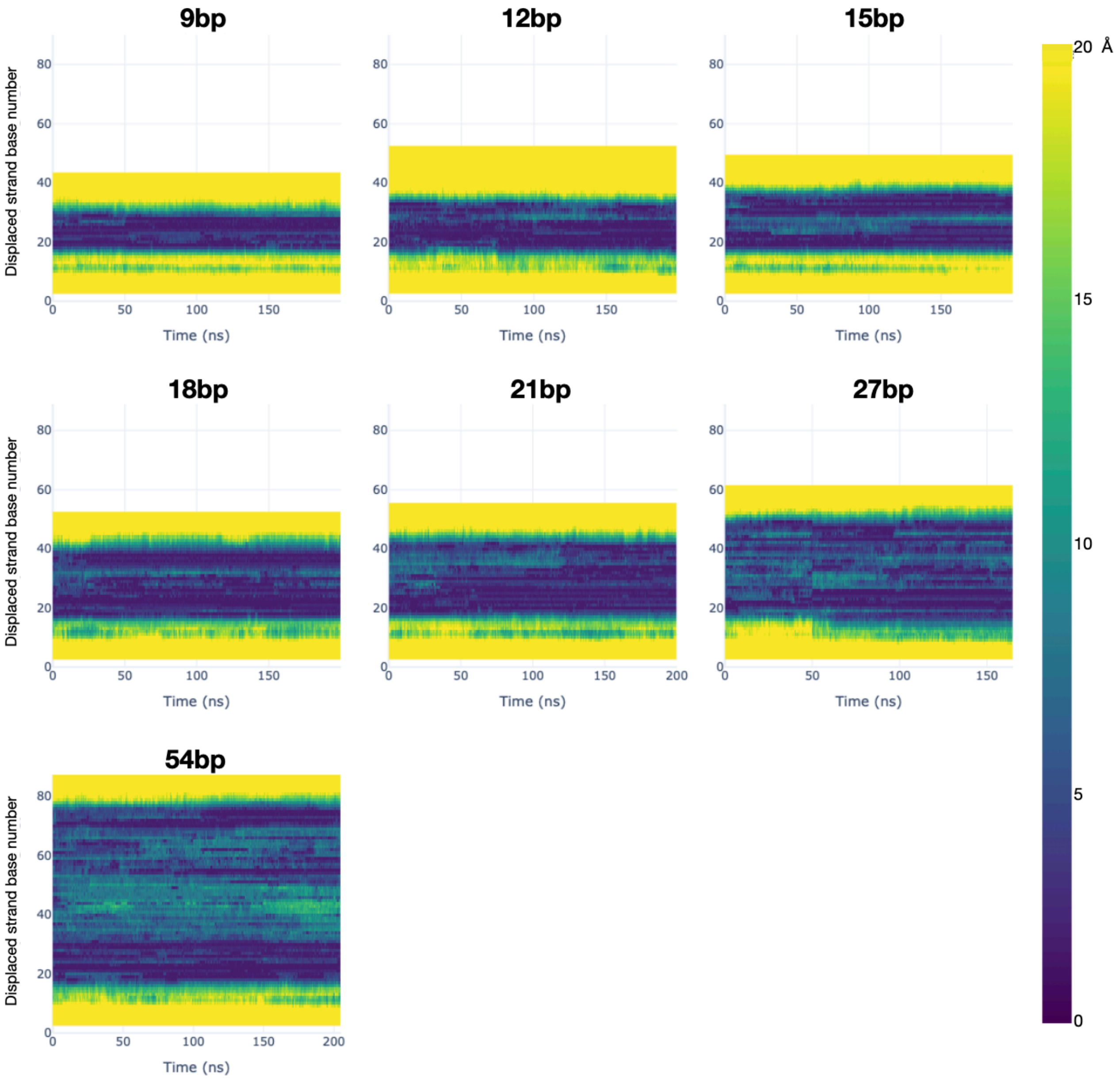


**Figure S8 -** Sequence dependent evolution of the distance from site II of the displaced strand phosphate groups

The y axis represents the numbering of the outgoing strand bases, from 5’ to 3’. Color shades indicate the proximity of the phosphate group of each base from site II along time, from dark blue to yellow.

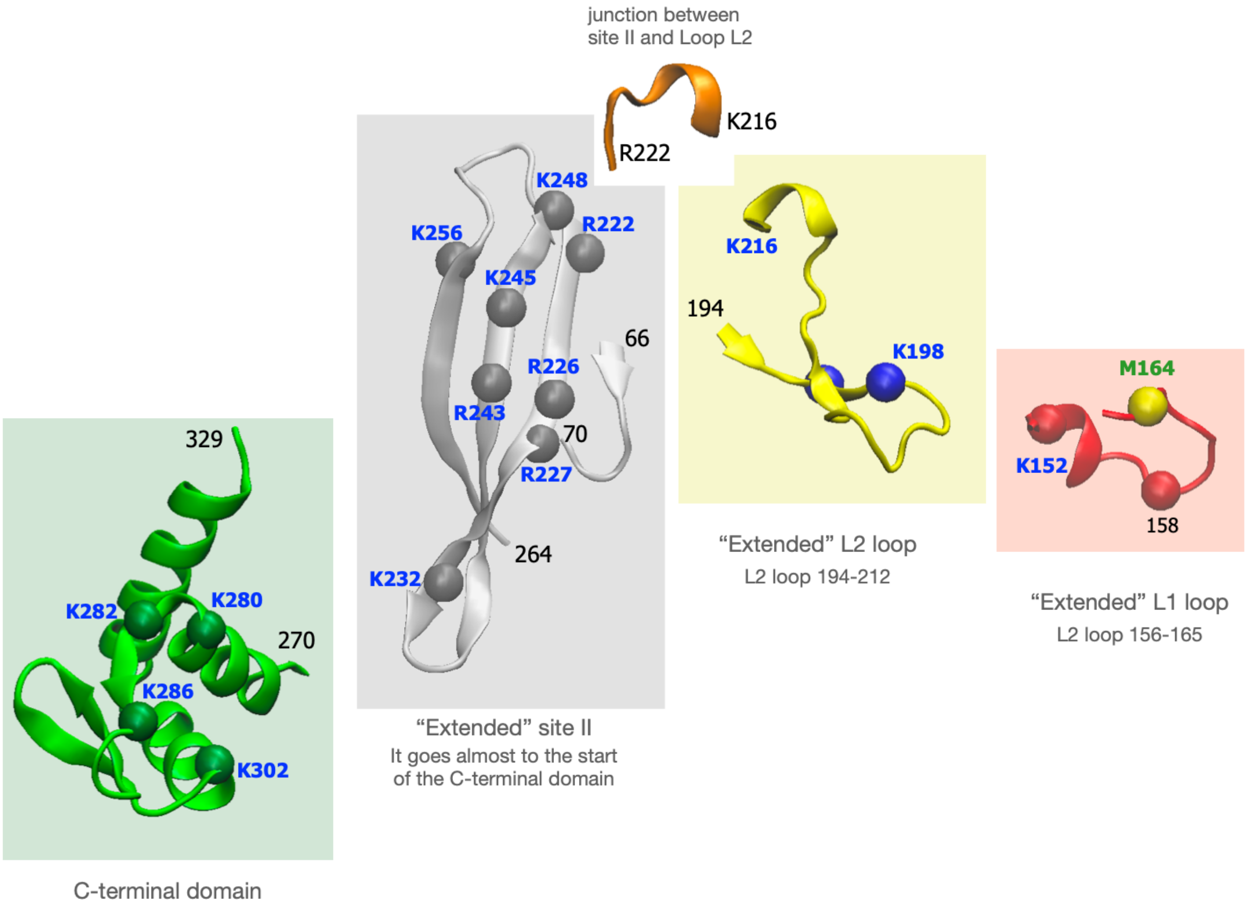


### Figure S9 - DNA binding motifs in the RecA nucleoprotein filaments of recombination

Split model of the DNA binding sites in RecA filaments. The C-terminal domain (green) non-specifically binds the incoming and outgoing dsDNA regions in B-form in 5' and in 3' of the D-loop. The "extended" site II (grey), that belongs to the rigid ATP-binding core domain, binds the D-loop region of the displaced strand; the "extended" L2 loop binds the complementary strand on its side that faces the L1 loop (site IV), and it can bind the displaced strand when that strand unbinds from site II on the loop side that faces site II (site III); the "extended" L1 loop binds the incoming strand in site I.

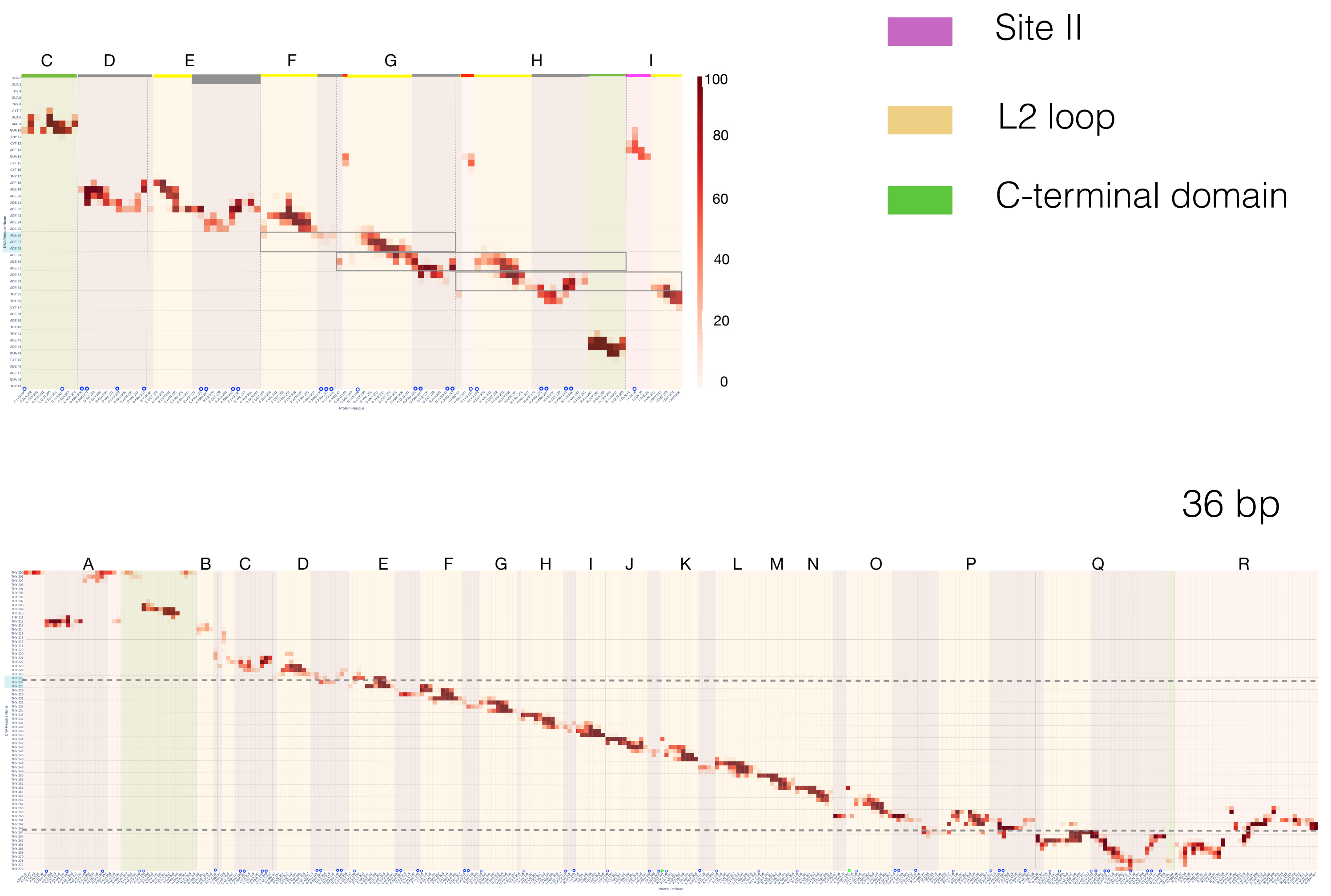


**Figure S10 - Contact maps for the 15bp and 36bp models**

The map displays the contacts of displaced strand phosphates (y axis) with residues of the RecA monomers, separated by vertical broken lines. Binding regions are shaded with different colors. The 15bp simulation shows a weakening of the site II binding to the dsDNA 5'-extremity also involves to a lesser extent two other monomers immediately in 5' of monomer *I*: in the contact map relative to the 15bp simulation displayed in supplementary, monomer *G* contacts the dsDNA via Glu158 and monomer *H* via Lys152.

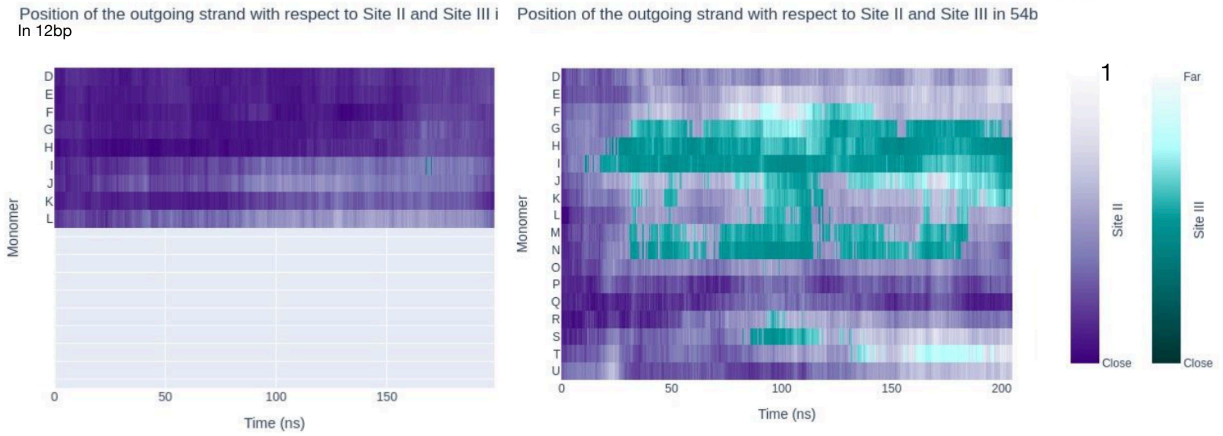


**Figure S11** - Site II versus site III occupancy during 200-ns MD simulation.

For each monomer (letters in the y-axis, from 5' to 3' in the alphabetic order), proximity of the outgoing strand phosphates to site II or site III is indicated using purple and green colors, respectively. The lowest value of the distance to site II or site III determines the color (values are normalized RMSDs, in Å).

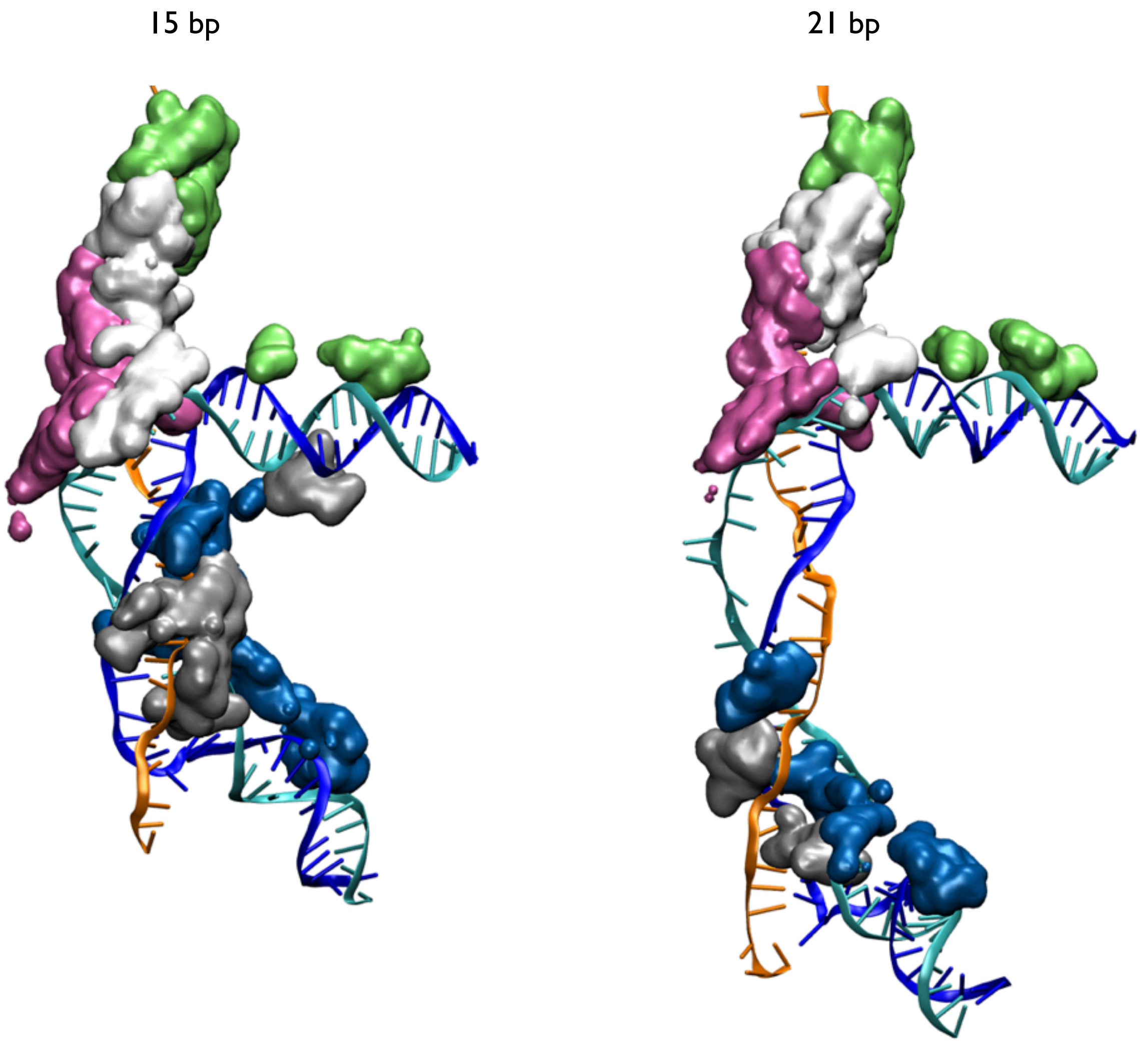


**Figure S12 -** D-loop binding regions of the filament monomers

Same as Figure 5, right insert, but protein regions that bind the ssDNA in 5' of the D-loop are also represented, in addition to the D-loop binding regions. Each protein is represented in a different color. The protein monomers that bind the upper side of the homoduplex B-form dsDNA extremity in 5' bridge that DNA region with the ssDNA in 5' but not with the D-loop.

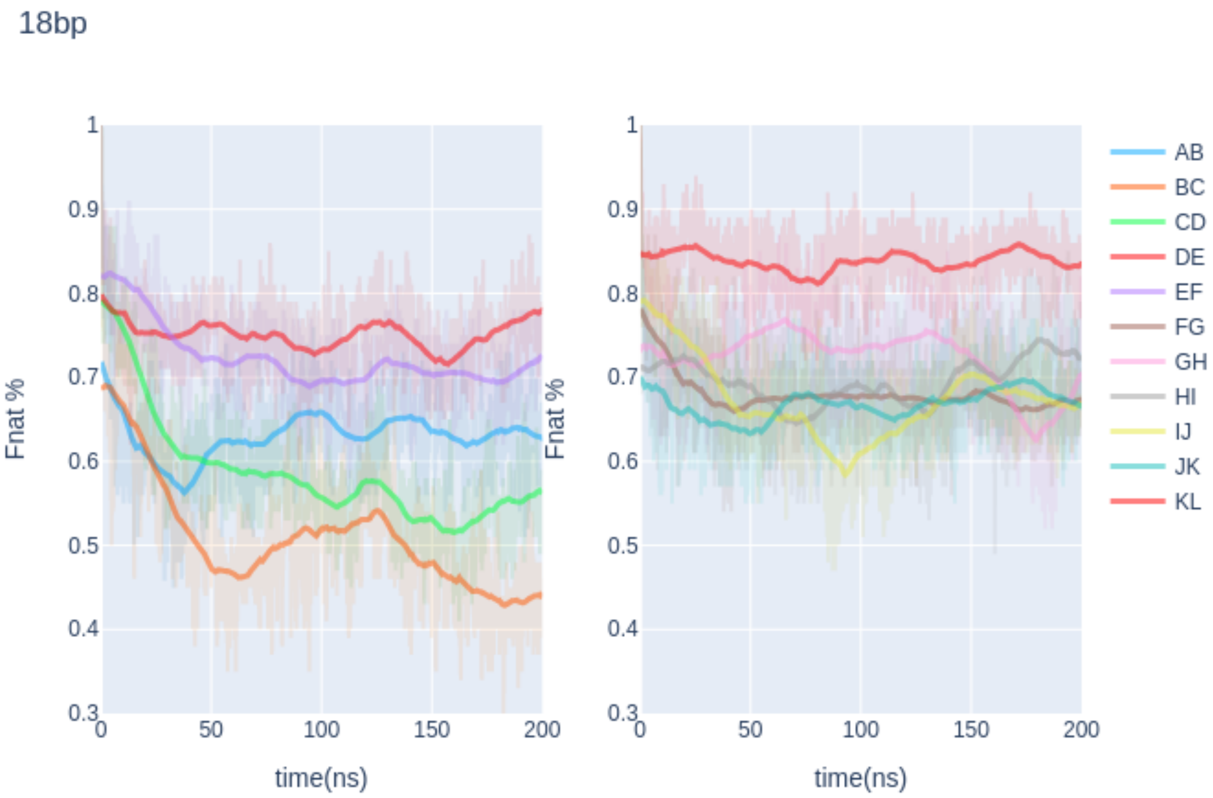


**Figure S13 -** Evolution of Fnat, the fraction of conserved interface contact pairs during 200ns of MD simulation of the 18bp model. Each interface is represented by a different color.
